# Deep learning detects virus presence in cancer histology

**DOI:** 10.1101/690206

**Authors:** Jakob Nikolas Kather, Jefree Schulte, Heike I. Grabsch, Chiara Loeffler, Hannah Muti, James Dolezal, Andrew Srisuwananukorn, Nishant Agrawal, Sara Kochanny, Saskia von Stillfried, Peter Boor, Takaki Yoshikawa, Dirk Jaeger, Christian Trautwein, Peter Bankhead, Nicole A. Cipriani, Tom Luedde, Alexander T. Pearson

**Affiliations:** Department of Medicine III, University Hospital RWTH Aachen, Aachen, Germany; German Cancer Consortium (DKTK), German Cancer Research Center (DKFZ), Heidelberg, Germany; Applied Tumor Immunity, German Cancer Research Center (DKFZ), Heidelberg, Germany; Department of Medicine, University of Chicago, Chicago, IL, USA; Department of Pathology, University of Chicago Medicine, Chicago, IL, USA; Pathology & Data Analytics, Leeds Institute of Medical Research at St James’s, University of Leeds, Leeds, UK; Department of Pathology, GROW School for Oncology and Developmental Biology, Maastricht University Medical Center+, Maastricht, The Netherlands; Department of Medicine, University of Illinois – Chicago, Chicago, IL, USA; Department of Surgery, University of Chicago, Chicago, IL, USA; Department of Pathology, University Hospital RWTH Aachen, Aachen, Germany; Department of Gastrointestinal Surgery, Kanagawa Cancer Center, Yokohama, Japan; Department of Gastric Surgery, National Cancer Center Hospital, Tokyo, Japan; MRC Institute of Genetics and Molecular Medicine, University of Edinburgh, Edinburgh, UK

## Abstract

Oncogenic viruses like human papilloma virus (HPV) or Epstein Barr virus (EBV) are a major cause of human cancer. Viral oncogenesis has a direct impact on treatment decisions because virus-associated tumors can demand a lower intensity of chemotherapy and radiation or can be more susceptible to immune check-point inhibition. However, molecular tests for HPV and EBV are not ubiquitously available.

We hypothesized that the histopathological features of virus-driven and non-virus driven cancers are sufficiently different to be detectable by artificial intelligence (AI) through deep learning-based analysis of images from routine hematoxylin and eosin (HE) stained slides. We show that deep transfer learning can predict presence of HPV in head and neck cancer with a patient-level 3-fold cross validated area-under-the-curve (AUC) of 0.89 [0.82; 0.94]. The same workflow was used for Epstein-Barr virus (EBV) driven gastric cancer achieving a cross-validated AUC of 0.80 [0.70; 0.92] and a similar performance in external validation sets. Reverse-engineering our deep neural networks, we show that the key morphological features can be made understandable to humans.

This workflow could enable a fast and low-cost method to identify virus-induced cancer in clinical trials or clinical routine. At the same time, our approach for feature visualization allows pathologists to look into the black box of deep learning, enabling them to check the plausibility of computer-based image classification.

## Introduction

Oncogenic viruses cause approximately 15% of malignant tumors in humans.^1^ Viruses can induce cancers with different histology and across different anatomic sites including squamous cell carcinomas (e.g. head and neck, cervix), adenocarcinomas (e.g. gastric), sarcomas (e.g. Kaposi), lymphomas (e.g. Burkitt) and hepatocellular carcinoma. Virus-driven tumors are an important health issue in western countries, but their global health impact is even higher as 80% of all virus-driven cancers occur in developing nations.^2^ Their incidence is expected to increase drastically in the next decade in developing and economically developed countries.^3,4^ Some types of cancer are almost always virally driven (e.g. cervical cancer) while others can have viral or non-viral driver mechanisms (e.g. head and neck cancer or gastric cancer). In these cases, it is important to determine if a patient’s tumor has a viral origin because if this is the case, a different clinical management may be warranted and virus status might influence the choice of a clinical trial for that particular patient. For example, in the case of head and neck squamous cell carcinoma (HNSC), patients with human papilloma virus (HPV)-positive tumors have superior overall survival compared to patients with HPV-negative tumors of the same stage and can benefit from treatment de-escalation.^5^ Likewise, patients with Epstein-Barr-Virus (EBV) related gastric adenocarcinoma tend to have a better prognosis and EBV positivity has been suggested as a biomarker for immunotherapy response.^6^

The gold standard method for detection of viruses in human cancer is dependent on the tumor type. In head and neck cancer, overexpression of p16 as assessed by immunohistochemistry is the most commonly used surrogate marker for virus presence. However, p16 is neither perfectly sensitive nor specific^7^, and some centers also use HPV polymerase-chain reaction, in-situ hybridization, or targeted DNA sequencing for HPV detection in tumor tissue. While these tests are more specific, they are also more expensive and time consuming. Presence of latent EBV infection in gastric cancer is usually measured using EBV-encoded RNA in-situ hybridization in pathology samples, which has a relatively high sensitivity and specificity but requires dedicated testing equipment and expertise for accurate interpretation.

In the present study, we hypothesized that morphological features correlating with the presence of viruses in solid tumors can be deduced from hematoxylin and eosin (H&E) histology, which is routinely available for almost any patient with a solid tumor. As a tool for feature extraction from images, we used deep learning, a form of artificial intelligence (AI), which has previously been used to detect high-level morphological features directly from histological images.^8-10^

## Methods

### Ethics and data sources

All experiments were conducted in accordance with the Declaration of Helsinki and the International Ethical Guidelines for Biomedical Research Involving Human Subjects. Anonymized scanned whole slide images were retrieved from The Cancer Genome Atlas (TCGA) project through the Genomics Data Commons Portal (https://portal.gdc.cancer.gov/). From this source, we retrieved images of head and neck squamous cell carcinoma (HNSC)^11^ and gastric adenocarcinoma (stomach adenocarcinoma, STAD)^12^. Exclusion criteria for patients in these cohorts were missing values in virus status, corrupt image files or lack of tumor tissue on the whole slide image. For TCGA-HNSC, images from N=450 patients were downloaded of which N=38 met exclusion criteria, leaving images from N=412 patients for further processing. For TCGA-STAD, images from N=416 patients were downloaded of which N=99 met exclusion criteria, leaving N=317 patients for further processing. Furthermore, we retrieved anonymized archival tissue samples of N=105 patients with HNSC from the University of Chicago Medicine Pathology archive (Chicago, Illinois, USA; “UCH-HNSC”) and anonymized tissue samples of N=197 patients with gastric cancer from the Kanagawa Cancer Center Hospital (Yokohama, Japan; “KCCH-STAD”) as described before^13^. For HNSC, HPV status was determined as described by Campbell et al.^14^ (by consensus of DNA sequencing^15^ and RNA sequencing^16^). For TCGA-STAD, EBV status was retrieved from genomic subtypes as described by Liu et al.^17^. For samples in UCH-HNSC, HPV status was defined by polymerase-chain reaction for the viral genes E6 and E7. For tumor samples in KCCH-STAD, EBV status was defined by EBV-encoded RNA in-situ hybridization.^18^

### Deep transfer learning workflow

All histological slides were reviewed and tumor regions were manually delineated in QuPath^19^, tessellated into tiles of 256 × 256 µm^2^ which were subsequently downsampled to 224 × 224 px, yielding an effective magnification of 1.14 µm/px. These tumor tiles were used for deep transfer learning in MATLAB R2019a as described before^9,10^. We used a modified VGG19 deep convolutional neural network^20^ which was pretrained on ImageNet (http://www.image-net.org, architecture shown in Suppl. Table 1). VGG19 was chosen because of its previously proven performance in detecting multiple tissue components in human cancer histology^9^ and because of its compatibility with the Deep Dream method (see below). All TCGA cohorts were randomly split into three equal subsets at patient level. A VGG19 classifier was trained on these data in a three-fold cross-validated way. This procedure yielded three independent classifiers which were evaluated on their respective test set of held-out patients. For each tumor type, the classifier was subsequently re-trained on the whole TCGA set and evaluated on an external test set.

### Feature visualization

To trace back deep-learning based predictions to human-understandable morphological patterns in histology, we used deep-dream-based visualization of output layer neurons for each class. We used a MATLAB implementation (https://de.mathworks.com/help/deeplearning/ref/deepdreamimage.html) of the original DeepDream algorithm (https://github.com/tensorflow/tensorflow/blob/master/tensorflow/examples/tutorials/deepdream/deepdream.ipynb) with pyramid level 6 and 500 iterations and subsequently auto-optimized colors by histogram stretching in IrfanView 4.52 (https://www.irfanview.com/). Color optimization was done with identical parameters for all deep-dream-images generated by a given network.

### Statistics and data presentation

Classifier performance was assessed by the Area under the Receiver Operating Curve (AUC under the ROC) with sensitivity (true positive rate, TPR) plotted on the vertical axis and 1 – specificity (false positive rate, FPR) plotted on the horizontal axis. 95% confidence intervals for the AUC were calculated with 500-fold bootstrapping with the “bias corrected and accelerated percentile method” ^28^ unless otherwise stated. For three-fold cross validated experiments, the mean of AUCs and the mean of confidence interval boundaries from all three classifiers is given if not otherwise noted. The ROC procedure is a widely used technique to assess the power of a classifier for any possible cutoff value of a numerical test. In this study, the cutoff for “percentage of virus-positive image tiles” was varied, yielding different sensitivity/specificity pairs which are plotted as ROC curves.

### Data availability

Images from the TCGA cohorts are available at https://portal.gdc.cancer.gov/. Our source codes are available at https://github.com/jnkather/VirusFromHE.

## Results

### Deep learning detects virus presence in squamous cell carcinomas and adenocarcinomas

We hypothesized that the presence of human papillomavirus (HPV) can be detected in head and neck squamous cell carcinoma (HNSC, Figure 1a) and that the presence Epstein-Barr-Virus (EBV) can be detected in gastric adenocarcinoma (STAD, Figure 1b) directly from histology by deep learning with a convolutional neural network (CNN). We used hematoxylin and eosin (H&E) stained tissue slides of patients included in the multicenter TCGA cohort (Suppl. Table 2) and trained a deep learning classifier in a patient-level three-fold cross-validated way (Figure 1c), followed by re-training on the whole cohort (Figure 1d). In head and neck cancer (N=412 patients, 12% HPV positive), this yielded an average patient-level AUC of 0.89 [0.82; 0.94] (Figure 1e) and applying the same workflow to detect EBV in gastric cancer (N=317 patients from TCGA, 8% EBV positive, Suppl. Table 3), a patient-level three-fold cross-validated neural network achieved an AUC of 0.80 [0.70; 0.92] (Figure 1f). Together, these results show that deep learning can robustly distinguish virus-induced (“virus present”) from non-virus-induced tumors (“virus not present”) across different histologies and anatomic sites.

**Figure 1:**
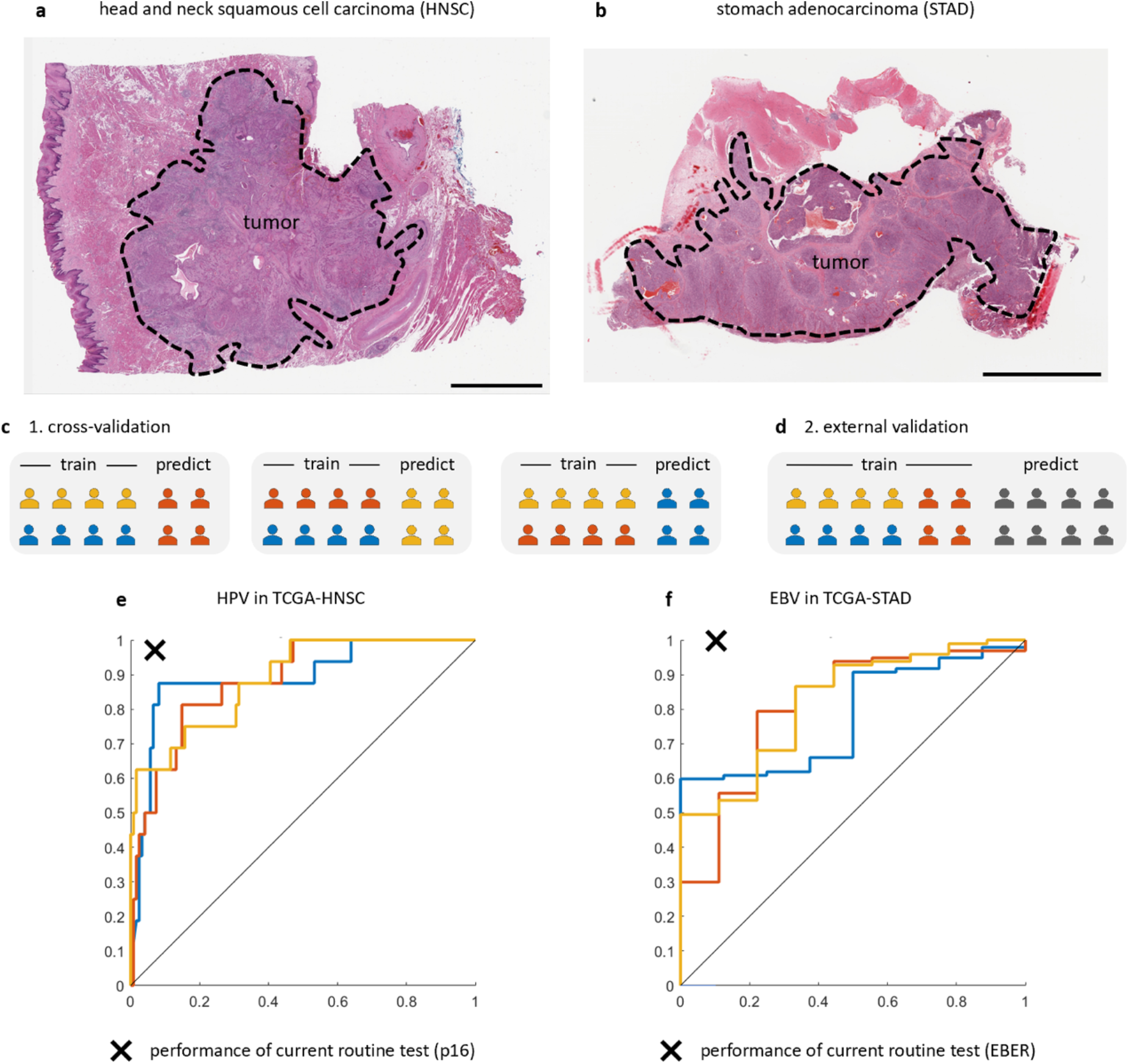
Detection of virus-induced cancer from histological routine images. (a) A representative sample (one of N=412 patients) of the TCGA-HNSC training cohort with a manual outline around the tumor tissue, (b) a representative sample of the TCGA-STAD cohort (1 of N=317 patients), scale bars in a and b are 5000 µm. (c) We trained and tested classifiers with patient-level three-fold cross-validation. (d) Subsequently, we re-trained on the whole cohort and tested in an external validation cohort. (e) Receiver operating characteristic (ROC, horizontal axis is 1-specificity and vertical axis is sensitivity) for HPV detection in TCGA-HNSC (N=412 patients), x marks the performance of the current clinical gold standard (p16 immunohistochemistry) (f) ROC for EBV detection in TCGA-STAD (N=317 patients), x marks the performance of EBV-encoded RNA (EBER) in-situ hybridization, the diagnostic gold standard. Each ROC curve corresponds to one cross-validation run.

### Noisy tile level data yields high patient-level accuracy in external validation cohorts

To assess the robustness of the classifiers, we used the neural network that was trained on the entire TCGA patient cohorts for head and neck and gastric cancer, respectively, and evaluated the classifiers on external validation cohorts. Non-overlapping tissue tiles of 256 µm edge length were used to predict a “virus probability score” which classified each tile as either virus positive or negative (derived from a tumor that was virally induced or derived from a tumor that was non-virally induced). These predictions were subsequently pooled on a patient level as “fraction of positive tiles” with varying thresholds according to the Receiver Operating Characteristic procedure (Figure 2a). Because each tile in the training set was assigned the label of the corresponding patient (obtained via bulk testing of tissue) and the tiles contained a multitude of different tissue types (tumor epithelium, stroma, necrosis, mucus, and others), this training set was inherently noisy. Correspondingly, predictions for virus-negative tiles in the EBV testing set were noisy with many false positive tile-level predictions (Figure 2b, right-hand side). However, tiles from virus-positive patients were mostly classified correctly (Figure 2a, left panel), enabling robust classification after pooling tile-level predictions on a patient level. AUC for EBV detection in the KCCH-STAD cohort (N=197 patients, 5% EBV positive) was 0.81 [0.69; 0.89] (Figure 2c; trained on TCGA-STAD, tested on KCCH-STAD). We manually assessed the histomorphology of tissue tiles in the KCCH-STAD cohort (Figure 2d, more examples are available at http://doi.org/10.5281/zenodo.3247009) and found that false positive tiles often presented with lymphocyte-rich stroma, a known morphological hallmark of EBV-positive gastric cancer.^21^ Thus, we conclude that misclassifications of individual image tiles were due to plausible human-understandable morphological features.

**Figure 2:**
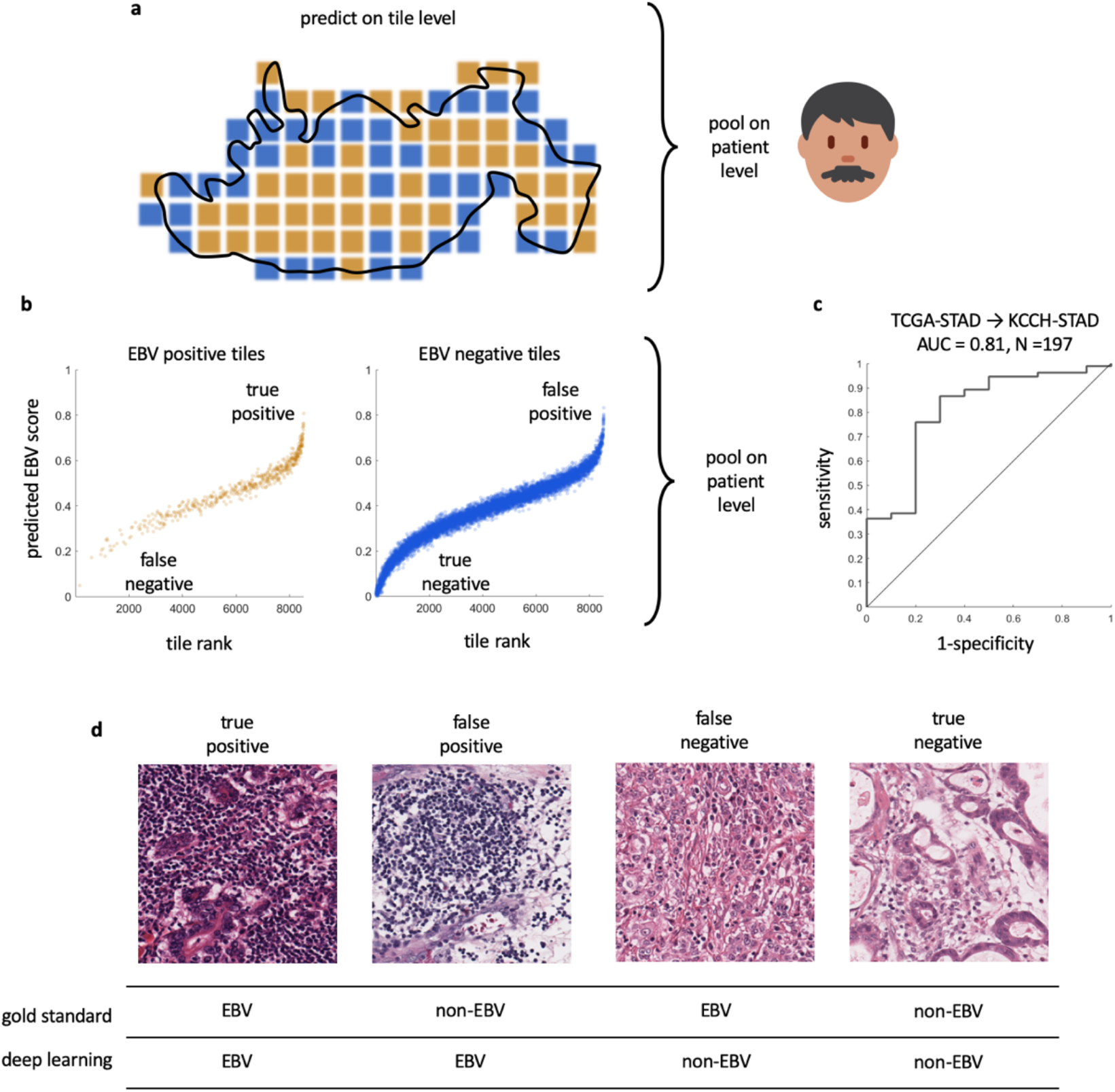
Tile-level classification yields high patient-level performance. (a) Schematic of the prediction process: in a histological whole slide image, non-overlapping areas of 256 x 256 µm (“blocks” or “tiles”) were used to predict virus status and subsequently pooled on a patient-level by fraction of positive tiles. Image credit for icon https://twemoji.twitter.com (b) For a subset of tiles in the KCCH-STAD validation cohort, the predicted EBV score is plotted for true EBV-positive (left) and EBV-negative tiles (right). A small random symmetric x-y-offset was added to each point for better visibility. While most positive tiles attained a high EBV score, prediction of EBV-negative tiles was noisy. More examples and more information about tile preprocessing is available at http://doi.org/10.5281/zenodo.3247009. (c) Despite noisy tile-level classification, patient-level prediction reached a high classification accuracy with an AUC of 0.81 in the independent KCCHSTAD validation set. (d) Representative tiles from the top and bottom quantile of EBV predictions. False positive tiles are lymphocyte-rich, which is a hallmark of virus-driven cancer and thus makes these misclassifications plausible.

Similarly, we validated the virus detector for HPV trained on TCGA-HNSC (N=412 patients, 12% HPV positive) in our in-house cohort UCH-HNSC from University of Chicago (N=105 patients, 49% HPV positive). This cohort had two main differences compared to the TCGA cohort which might negatively affect classifier performance: first, a polymerase-chain reaction for high-risk HPV viral genes was used to determine virus status. Second, this cohort was artificially balanced for HPV status and thus had a much higher prevalence of HPV-induced cancer than TCGA. In spite of these stark differences, our classifier achieved an AUC of 0.70 [0.66; 0.74] for HPV prediction in UCH-HNSC. Manual review of representative tissue tiles by an expert pathologist showed that tiles with a high HPV prediction score were “carcinomas with large nested, broad-based invasion and relative decrease in cytoplasmic keratinization, resulting in a blue (cooltoned) appearance”. This is compatible with previously known morphological features of HPV-positive HNSC.^22^ Tiles with a low HPV prediction score were “carcinomas with small nested invasion and eosinophilic (pink or warm-toned) cytoplasmic keratinization”. Thus, we conclude that in HNSC as well as in gastric cancer, predictions of viral status in individual tissue tiles by a deep neural network were plausible to expert pathologists.

Together, these data show that despite noisy training data and tile-level misclassifications, patient-level prediction of virus status in HNSC and gastric cancer can reach a high accuracy.

### Reverse-engineering trained neural networks

Attempting to characterize more precisely which morphological features may have been used by the neuronal network to detect virus-induced cancer, we used a feature-visualization method and discussed the results with a panel of expert pathologists. We hypothesized that reverse-engineering features from neural networks could be used as a plausibility check for deep learning, completing the cycle “human to AI and back”. In an exploratory study, we employed the Deep Dream algorithm which uses a trained neural network (Figure 3a) to create pseudo-images for each output class in the classification layer (Figure 3b). This approach yielded “pseudo-histology” images for HPV positive and negative HNSC (Figure 3b) and EBV positive and negative gastric cancer (Figure 3c), discussing the resulting images with five pathologists. In general, pathologists described the images as “beautiful” and “psychedelic”. Relating the aspect of pseudo-histology in histological terms, they described the features as “a sheet of small nodules composed of bright, predominantly warm colors” (HPV negative HNSC, Figure 3c, left panel), “large nests with rounded borders composed of dark, predominantly cool colors punctuated by red dots” (HPV positive HNSC, Figure 3c, right panel) and “ill-defined dark green whorls punctuated by blue dots and wisps of yellow” (EBV negative gastric cancer [STAD], Figure 3d, left panel), “overlapping sheets with reticulated patterns in pastel colors”, potentially resembling “lymphoid stroma” (EBV positive gastric cancer, Figure 3d right panel).

**Figure 3:**
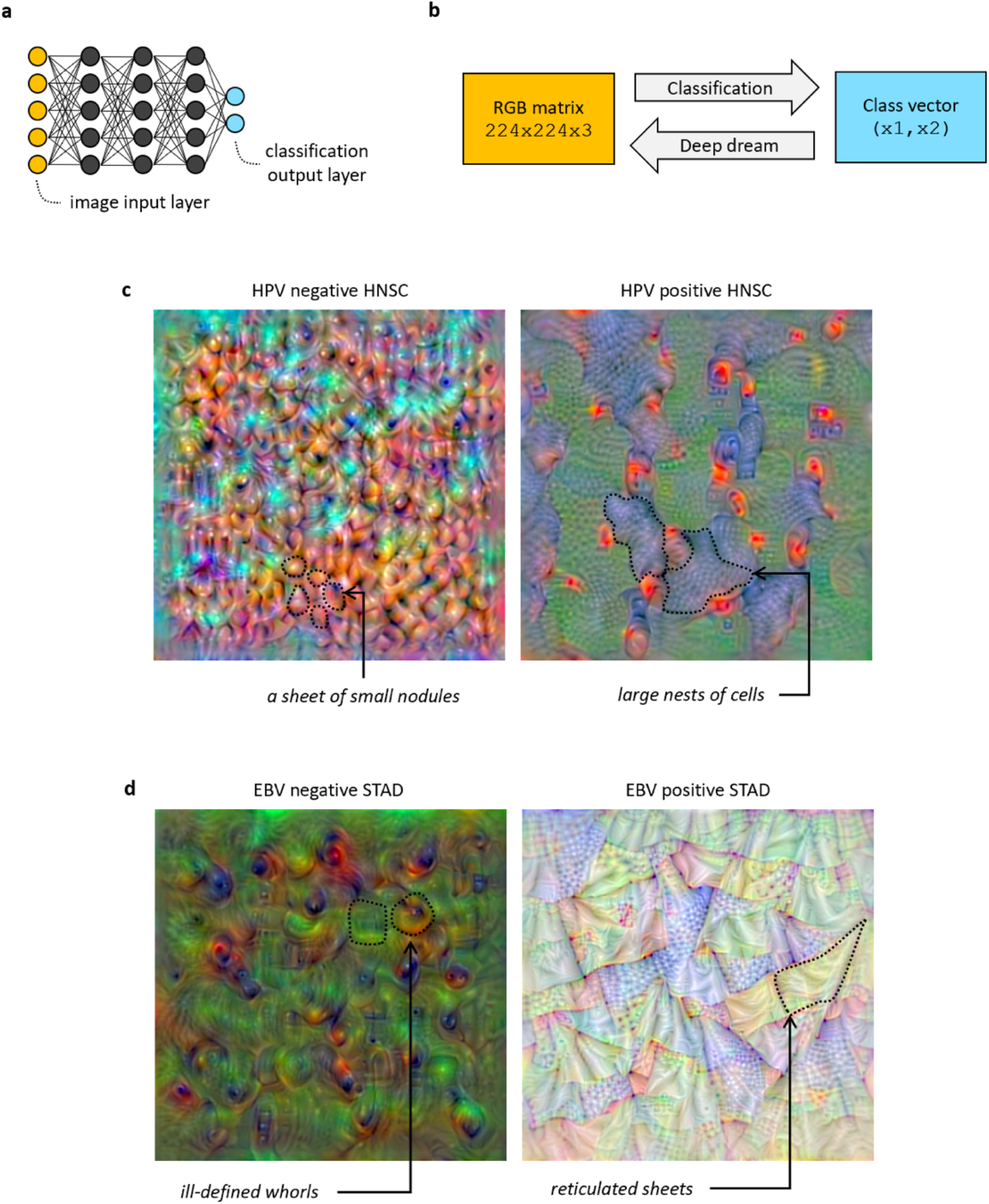
Feature visualization of viral morphological signatures in histological images by Deep Dream. (a) We used a modified VGG19 deep neural network that reads images in through the input layer and outputs predictions in a two-neuron output layer. (b) Information flow from left to right is used to classify images. The Deep dream algorithm uses the reverse direction to iteratively create pseudo images for each output neuron. (c) Example of Deep dream pseudo-images for HPV negative and positive HNSC with subjective manual description on morphological features by expert pathologists, (b) corresponding images for EBV in STAD.

Together, these data show that deep learning can plausibly sort tissue tiles (Figure 2d) and yields a high classification performance for virus presence (Figure 2c). The actual morphological patterns used for this classification may be different from the ones that humans typically use but can be visualized in a way that might be understandable for humans through the Deep Dream algorithm (as has been shown in nonmedical applications^23^). Based on this, we conclude that, by analyzing tile-level classification and possibly by analyzing Deep Dream images, human observers can get an insight about morphological patterns used by deep neural networks, allowing for quality control and possibly performing human classification performance through machine-identified morphological features.

## Discussion

Virus-induced cancers mostly occur in developing countries, making them neglected diseases on a global scale. Some of these virus-driven cancers are under-tested for in clinical routine. Existing wet lab assays to test for virus presence (such as sequencing) are costly, require a high level of expertise (such as in-situ hybridization) and not all assays achieve a perfect classification accuracy (such as p16 immunohistochemistry^24^). Here, we present a deep-learning-based low-to-no-cost assay for routine detection of virus presence from ubiquitously available histology in two major tumor types of very different histology. We demonstrate that classification accuracy is as high as AUC 0.81 when trained with a few hundred patients. Our approach relies on digitally scanned images of hematoxylin & eosin stained tissue slides. The cost to scan such a histology slide is well below $10 at low throughput and considerably lower at high throughput^10^, potentially enabling noticeable cost savings for virus testing of tumor tissue in the future.

At the moment, sensitivity and specificity of our classifier is lower than in routine diagnostic tests: for EBV detection in gastric cancer by EBV-encoded RNA in-situ hybridization, one study reported a sensitivity of 100% at a specificity of 90%.^25^ For HPV detection in HNSC by p16 immunohistochemistry, another study reported a sensitivity of 97.4% and a specificity 93.75%.^24^ As shown in Figure 1e-f and Figure 2c, the deep learning classifier approaches these gold standard methods but is still less sensitive and specific. However, sensitivity and specificity of our method are higher than those in previous studies of deep-learning based prediction of molecular features from histology.^8,26^

In our experiments, the deep learning classifiers reached a high cross-validated performance which we could replicate in an external validation set for gastric cancer (Figure 2c). In the multicenter TCGA-HNSC cohort, cross-validated HPV detection performance was high, but dropped in the external validation cohort UCH-HNSC. This may be related to the relatively small patient size of this cohort or due to different gold standards for HPV detection (consensus of DNA and RNA sequencing in TCGA-HNSC and polymerase-chain reaction in UCH-HNSC). Most probably, however, this is due to the very different prevalence of virus-induced cancers in the training set and in the test set. Whereas the training set (TCGA-HNSC) reflected the natural prevalence of HPV-positive cancers, the test set (UCH-HNSC) was artificially balanced to a prevalence of 50%. This may have negatively affected classifier performance as has been described for mutation prediction in lung cancer^8^.

According to our experience from similar tasks, it can be expected that training on larger clinical cohorts will likely improve performance of our method. Similarly, further optimizing hyperparameters and neural network architectures will likely yield a performance boost. Also, further dividing deep learning classifiers by anatomical sub-sites of tumors (e.g. oropharyngeal or hypopharyngeal) will likely increase performance. In the end, this image-based biomarker, like all biomarkers, needs to be tested in prospective clinical trials before widespread clinical use.

A new aspect of our study is the approach “human to AI and back”: humans (expert pathologists) delineated tumor tissue in whole slide sections and thus enabled the AI to detect virus presence in histological images. In turn, using deep-dream-based feature visualization, we show that the AI can in principle inform a human observer about morphological features of interest. Feature visualization by Deep Dream and similar methods^27^ is well-established to understand the inner workings of deep neural networks. Yet, to our knowledge, this has never been systematically used for pathologist-AI-crosstalk. Thus, our study shows for the first time that deep learning algorithms can not only be used as tools to facilitate diagnostic routine but could also enable human observers to get a different viewpoint on histomorphology.

## Acknowledgements

The results are in part based upon data generated by the TCGA Research Network: http://cancergenome.nih.gov/. Our funding sources are as follows. J.N.K.: RWTH University Aachen (START 2018-691906). S. v. S.: RWTH University Aachen (START 2017-691742) and DFG (GRK 2375/1). P. Boor: DFG (SFB-TRR57/P06, P25, M01, SFB-TRR219, LU 1360/3-1, BO 3755/3-1 and 6-1), BMBF (01GM1901A) and IZKF (O3-7). T.L.: Horizon 2020 through the European Research Council (ERC) Consolidator Grant PhaseControl (771083), Mildred-Scheel-Endowed Professorship from the German Cancer Aid, DFG (SFB-TRR57/P06, LU 1360/3-1), Ernst-Jung-Foundation Hamburg and IZKF (interdisciplinary center of clinical research) at RWTH Aachen. A.T.P.: NIH/NIDCR (#K08-DE026500), Institutional Research Grant (#IRG-16-222-56) from the American Cancer Society, and the University of Chicago Medicine Comprehensive Cancer Center Support Grant (#P30-CA14599). The authors would also like to thank the generous support from Chef Grant Achatz, Chef Giuseppe Tentori, and Nick Kokonas.

