## Supplementary material for "Deep learning detects virus presence in cancer histology": Suppl. Tables 1-3

### Supplementary data

| | Layer Name | Layer Type | Layer Description |
|---|---|---|---|
| **1** | 'input' | Image Input | 224x224x3 images with 'zerocenter' normalization |
| **2** | 'conv1_1' | Convolution | 64 3x3x3 convolutions with stride [1 1] and padding [1 1 1 1] |
| **3** | 'relu1_1' | ReLU | ReLU |
| **4** | 'conv1_2' | Convolution | 64 3x3x64 convolutions with stride [1 1] and padding [1 1 1 1] |
| **5** | 'relu1_2' | ReLU | ReLU |
| **6** | 'pool1' | Max Pooling | 2x2 max pooling with stride [2 2] and padding [0 0 0 0] |
| **7** | 'conv2_1' | Convolution | 128 3x3x64 convolutions with stride [1 1] and padding [1 1 1 1] |
| **8** | 'relu2_1' | ReLU | ReLU |
| **9** | 'conv2_2' | Convolution | 128 3x3x128 convolutions with stride [1 1] and padding [1 1 1 1] |
| **10** | 'relu2_2' | ReLU | ReLU |
| **11** | 'pool2' | Max Pooling | 2x2 max pooling with stride [2 2] and padding [0 0 0 0] |
| **12** | 'conv3_1' | Convolution | 256 3x3x128 convolutions with stride [1 1] and padding [1 1 1 1] |
| **13** | 'relu3_1' | ReLU | ReLU |
| **14** | 'conv3_2' | Convolution | 256 3x3x256 convolutions with stride [1 1] and padding [1 1 1 1] |
| **15** | 'relu3_2' | ReLU | ReLU |
| **16** | 'conv3_3' | Convolution | 256 3x3x256 convolutions with stride [1 1] and padding [1 1 1 1] |
| **17** | 'relu3_3' | ReLU | ReLU |
| **18** | 'conv3_4' | Convolution | 256 3x3x256 convolutions with stride [1 1] and padding [1 1 1 1] |
| **19** | 'relu3_4' | ReLU | ReLU |
| **20** | 'pool3' | Max Pooling | 2x2 max pooling with stride [2 2] and padding [0 0 0 0] |
| **21** | 'conv4_1' | Convolution | 512 3x3x256 convolutions with stride [1 1] and padding [1 1 1 1] |
| **22** | 'relu4_1' | ReLU | ReLU |
| **23** | 'conv4_2' | Convolution | 512 3x3x512 convolutions with stride [1 1] and padding [1 1 1 1] |
| **24** | 'relu4_2' | ReLU | ReLU |

| | | | |
|---|---|---|---|
| **25** | 'conv4_3' | Convolution | 512 3x3x512 convolutions with stride [1 1] and padding [1 1 1 1] |
| **26** | 'relu4_3' | ReLU | ReLU |
| **27** | 'conv4_4' | Convolution | 512 3x3x512 convolutions with stride [1 1] and padding [1 1 1 1] |
| **28** | 'relu4_4' | ReLU | ReLU |
| **29** | 'pool4' | Max Pooling | 2x2 max pooling with stride [2 2] and padding [0 0 0 0] |
| **30** | 'conv5_1' | Convolution | 512 3x3x512 convolutions with stride [1 1] and padding [1 1 1 1] |
| **31** | 'relu5_1' | ReLU | ReLU |
| **32** | 'conv5_2' | Convolution | 512 3x3x512 convolutions with stride [1 1] and padding [1 1 1 1] |
| **33** | 'relu5_2' | ReLU | ReLU |
| **34** | 'conv5_3' | Convolution | 512 3x3x512 convolutions with stride [1 1] and padding [1 1 1 1] |
| **35** | 'relu5_3' | ReLU | ReLU |
| **36** | 'conv5_4' | Convolution | 512 3x3x512 convolutions with stride [1 1] and padding [1 1 1 1] |
| **37** | 'relu5_4' | ReLU | ReLU |
| **38** | 'pool5' | Max Pooling | 2x2 max pooling with stride [2 2] and padding [0 0 0 0] |
| **39** | 'fc6' | Fully Connected | 4096 fully connected layer |
| **40** | 'relu6' | ReLU | ReLU |
| **41** | 'drop6' | Dropout | 50% dropout |
| **42** | 'fc7' | Fully Connected | 4096 fully connected layer |
| **43** | 'relu7' | ReLU | ReLU |
| **44** | 'drop7' | Dropout | 50% dropout |
| **45** | 'fc' | Fully Connected | 2 fully connected layer |
| **46** | 'prob' | Softmax | softmax |
| **47** | 'classoutput' | Classification Output | crossentropyex with classes 'negative' and 'positive' |

**Suppl. Table 1: Transfer learning network derived from VGG19**

| Patient | Sex | Stage | Site of origin | HPV status |
|---|---|---|---|---|
| **TCGA-4P-AA8J** | male | stage iva | Tongue, NOS | negative |
| **TCGA-BA-4076** | male | N/A | Larynx, NOS | negative |
| **TCGA-BA-4078** | male | N/A | Larynx, NOS | negative |
| **TCGA-BA-5149** | male | stage iva | Floor of mouth, NOS | negative |
| **TCGA-BA-5151** | male | stage iva | Cheek mucosa | negative |
| **TCGA-BA-5152** | male | stage iva | Overlapping lesion of lip, oral cavity and pharynx | negative |
| **TCGA-BA-5153** | male | N/A | Tonsil, NOS | positive |
| **TCGA-BA-5555** | male | stage iva | Larynx, NOS | negative |
| **TCGA-BA-5556** | female | stage ii | Floor of mouth, NOS | negative |
| **TCGA-BA-5558** | male | N/A | Overlapping lesion of lip, oral cavity and pharynx | negative |
| **TCGA-BA-5559** | male | N/A | Tonsil, NOS | positive |
| **TCGA-BA-6868** | male | N/A | Larynx, NOS | negative |
| **TCGA-BA-6869** | male | stage iii | Larynx, NOS | positive |
| **TCGA-BA-6871** | male | N/A | Base of tongue, NOS | negative |
| **TCGA-BA-6872** | male | N/A | Floor of mouth, NOS | negative |
| **TCGA-BA-6873** | male | stage iva | Tongue, NOS | negative |
| **TCGA-BA-A6D8** | male | stage iva | Floor of mouth, NOS | negative |
| **TCGA-BA-A6DA** | female | stage iva | Supraglottis | negative |
| **TCGA-BA-A6DB** | female | stage i | Tongue, NOS | negative |
| **TCGA-BA-A6DD** | male | stage iva | Floor of mouth, NOS | negative |
| **TCGA-BA-A6DE** | female | stage ii | Tongue, NOS | negative |
| **TCGA-BA-A6DG** | male | N/A | Tongue, NOS | negative |
| **TCGA-BA-A6DI** | male | N/A | Larynx, NOS | negative |
| **TCGA-BA-A6DJ** | male | stage iva | Gum, NOS | negative |
| **TCGA-BA-A6DL** | male | N/A | Oropharynx, NOS | negative |
| **TCGA-BA-A8YP** | male | stage ivb | Oropharynx, NOS | negative |

| | | | | |
|---|---|---|---|---|
| **TCGA-BB-4217** | male | stage iva | Larynx, NOS | negative |
| **TCGA-BB-4223** | male | stage iva | Tonsil, NOS | positive |
| **TCGA-BB-4224** | male | stage iva | Tongue, NOS | negative |
| **TCGA-BB-4228** | male | stage iii | Base of tongue, NOS | positive |
| **TCGA-BB-7861** | male | stage iii | Base of tongue, NOS | positive |
| **TCGA-BB-7862** | male | stage iva | Larynx, NOS | positive |
| **TCGA-BB-7863** | female | stage iii | Tongue, NOS | negative |
| **TCGA-BB-7864** | male | stage iva | Larynx, NOS | positive |
| **TCGA-BB-7866** | male | N/A | Tonsil, NOS | positive |
| **TCGA-BB-7870** | male | stage iva | Larynx, NOS | negative |
| **TCGA-BB-7872** | male | stage i | Mouth, NOS | negative |
| **TCGA-BB-8596** | female | stage iva | Hypopharynx, NOS | negative |
| **TCGA-BB-8601** | male | stage iii | Floor of mouth, NOS | negative |
| **TCGA-BB-A5HU** | male | stage iva | Mouth, NOS | negative |
| **TCGA-BB-A5HY** | male | stage iva | Hypopharynx, NOS | negative |
| **TCGA-BB-A5HZ** | male | stage iva | Mouth, NOS | negative |
| **TCGA-BB-A6UM** | male | stage iii | Tonsil, NOS | positive |
| **TCGA-BB-A6UO** | female | stage iva | Tongue, NOS | negative |
| **TCGA-C9-A47Z** | female | stage iii | Tongue, NOS | negative |
| **TCGA-C9-A480** | female | stage iii | Tongue, NOS | negative |
| **TCGA-CN-4722** | female | stage ii | Larynx, NOS | negative |
| **TCGA-CN-4723** | male | stage iva | Larynx, NOS | negative |
| **TCGA-CN-4725** | male | stage ii | Tongue, NOS | negative |
| **TCGA-CN-4726** | male | stage iva | Cheek mucosa | negative |
| **TCGA-CN-4727** | male | stage iva | Larynx, NOS | negative |
| **TCGA-CN-4728** | male | stage iva | Overlapping lesion of lip, oral cavity and pharynx | negative |
| **TCGA-CN-4729** | male | stage iii | Overlapping lesion of lip, oral cavity and pharynx | negative |

| | | | | |
|---|---|---|---|---|
| **TCGA-CN-4730** | male | stage iva | Overlapping lesion of lip, oral cavity and pharynx | negative |
| **TCGA-CN-4731** | female | stage iva | Overlapping lesion of lip, oral cavity and pharynx | negative |
| **TCGA-CN-4733** | male | stage iii | Tongue, NOS | negative |
| **TCGA-CN-4734** | male | stage ii | Cheek mucosa | negative |
| **TCGA-CN-4735** | male | stage iva | Larynx, NOS | negative |
| **TCGA-CN-4736** | female | N/A | Tongue, NOS | negative |
| **TCGA-CN-4738** | male | stage iva | Larynx, NOS | negative |
| **TCGA-CN-4741** | male | stage iva | Gum, NOS | positive |
| **TCGA-CN-4742** | female | stage iva | Tongue, NOS | negative |
| **TCGA-CN-5355** | male | stage iva | Larynx, NOS | negative |
| **TCGA-CN-5356** | male | stage iii | Larynx, NOS | negative |
| **TCGA-CN-5358** | male | stage ii | Floor of mouth, NOS | negative |
| **TCGA-CN-5359** | male | stage iva | Floor of mouth, NOS | negative |
| **TCGA-CN-5360** | male | stage iva | Larynx, NOS | negative |
| **TCGA-CN-5361** | male | stage iva | Larynx, NOS | negative |
| **TCGA-CN-5363** | male | stage ivb | Larynx, NOS | negative |
| **TCGA-CN-5364** | male | stage iva | Floor of mouth, NOS | negative |
| **TCGA-CN-5365** | male | stage ivb | Tonsil, NOS | negative |
| **TCGA-CN-5366** | male | stage iva | Hypopharynx, NOS | negative |
| **TCGA-CN-5367** | female | stage iva | Tongue, NOS | negative |
| **TCGA-CN-5369** | female | stage iva | Hard palate | negative |
| **TCGA-CN-5373** | female | stage i | Floor of mouth, NOS | negative |
| **TCGA-CN-5374** | female | stage iva | Tonsil, NOS | positive |
| **TCGA-CN-6010** | male | stage iva | Larynx, NOS | negative |
| **TCGA-CN-6012** | male | stage iva | Larynx, NOS | negative |
| **TCGA-CN-6013** | male | stage iva | Gum, NOS | negative |
| **TCGA-CN-6016** | male | stage iva | Tongue, NOS | negative |

| | | | | |
|---|---|---|---|---|
| **TCGA-CN-6017** | male | stage iva | Tongue, NOS | negative |
| **TCGA-CN-6018** | female | stage iva | Overlapping lesion of lip, oral cavity and pharynx | negative |
| **TCGA-CN-6019** | male | stage iva | Tongue, NOS | negative |
| **TCGA-CN-6021** | female | N/A | Larynx, NOS | negative |
| **TCGA-CN-6022** | male | stage iva | Larynx, NOS | negative |
| **TCGA-CN-6023** | male | stage iva | Larynx, NOS | negative |
| **TCGA-CN-6024** | male | stage iva | Tongue, NOS | negative |
| **TCGA-CN-6988** | male | stage iva | Larynx, NOS | negative |
| **TCGA-CN-6989** | male | stage iva | Larynx, NOS | negative |
| **TCGA-CN-6992** | male | stage iva | Larynx, NOS | negative |
| **TCGA-CN-6994** | male | stage iva | Overlapping lesion of lip, oral cavity and pharynx | negative |
| **TCGA-CN-6996** | female | stage iva | Tongue, NOS | negative |
| **TCGA-CN-6997** | male | stage iva | Larynx, NOS | negative |
| **TCGA-CN-A63T** | male | stage iva | Larynx, NOS | negative |
| **TCGA-CN-A63U** | male | stage iva | Larynx, NOS | negative |
| **TCGA-CN-A63V** | male | stage iva | Cheek mucosa | negative |
| **TCGA-CN-A63W** | female | stage iva | Larynx, NOS | negative |
| **TCGA-CN-A641** | male | stage iva | Larynx, NOS | negative |
| **TCGA-CN-A642** | male | stage ivb | Tongue, NOS | negative |
| **TCGA-CQ-5323** | male | stage i | Overlapping lesion of lip, oral cavity and pharynx | positive |
| **TCGA-CQ-5324** | male | stage iii | Overlapping lesion of lip, oral cavity and pharynx | negative |
| **TCGA-CQ-5325** | male | stage i | Tongue, NOS | negative |
| **TCGA-CQ-5326** | male | stage iva | Overlapping lesion of lip, oral cavity and pharynx | negative |
| **TCGA-CQ-5329** | female | stage ii | Tongue, NOS | negative |
| **TCGA-CQ-5330** | female | stage iva | Tongue, NOS | negative |
| **TCGA-CQ-5331** | female | N/A | Overlapping lesion of lip, oral cavity and pharynx | negative |
| **TCGA-CQ-5332** | male | stage iii | Overlapping lesion of lip, oral cavity and pharynx | negative |

| | | | | |
|---|---|---|---|---|
| **TCGA-CQ-5333** | male | stage ii | Tongue, NOS | negative |
| **TCGA-CQ-5334** | male | stage iva | Cheek mucosa | negative |
| **TCGA-CQ-6218** | female | stage iva | Tongue, NOS | negative |
| **TCGA-CQ-6219** | female | stage iva | Tongue, NOS | negative |
| **TCGA-CQ-6220** | male | stage iii | Overlapping lesion of lip, oral cavity and pharynx | negative |
| **TCGA-CQ-6222** | male | stage iva | Tongue, NOS | negative |
| **TCGA-CQ-6223** | male | stage ii | Overlapping lesion of lip, oral cavity and pharynx | negative |
| **TCGA-CQ-6224** | male | stage iva | Tongue, NOS | negative |
| **TCGA-CQ-6225** | male | stage iii | Overlapping lesion of lip, oral cavity and pharynx | negative |
| **TCGA-CQ-6227** | male | stage iva | Overlapping lesion of lip, oral cavity and pharynx | negative |
| **TCGA-CQ-6228** | female | stage iva | Overlapping lesion of lip, oral cavity and pharynx | negative |
| **TCGA-CQ-6229** | male | stage ii | Tongue, NOS | negative |
| **TCGA-CQ-7063** | female | stage i | Mouth, NOS | negative |
| **TCGA-CQ-7065** | male | stage ii | Tongue, NOS | negative |
| **TCGA-CQ-7067** | female | stage i | Mouth, NOS | negative |
| **TCGA-CQ-7068** | female | stage ii | Overlapping lesion of lip, oral cavity and pharynx | negative |
| **TCGA-CQ-7069** | female | stage ii | Mouth, NOS | negative |
| **TCGA-CQ-7071** | female | stage iii | Mouth, NOS | negative |
| **TCGA-CQ-7072** | male | stage ii | Tongue, NOS | negative |
| **TCGA-CQ-A4C6** | male | stage ii | Cheek mucosa | negative |
| **TCGA-CQ-A4C9** | male | stage iii | Anterior floor of mouth | negative |
| **TCGA-CQ-A4CA** | male | stage ii | Ventral surface of tongue, NOS | negative |
| **TCGA-CQ-A4CB** | male | stage iii | Floor of mouth, NOS | negative |
| **TCGA-CQ-A4CD** | male | stage iva | Mandible | negative |
| **TCGA-CQ-A4CE** | female | stage ii | Tongue, NOS | negative |
| **TCGA-CQ-A4CG** | female | stage iii | Cheek mucosa | negative |
| **TCGA-CQ-A4CH** | male | stage ii | Border of tongue | negative |

| | | | | |
|---|---|---|---|---|
| **TCGA-CQ-A4CI** | male | stage iii | Cheek mucosa | negative |
| **TCGA-CV-5430** | male | stage iva | Larynx, NOS | negative |
| **TCGA-CV-5431** | male | stage iva | Larynx, NOS | negative |
| **TCGA-CV-5432** | male | stage iii | Larynx, NOS | negative |
| **TCGA-CV-5434** | male | stage iva | Larynx, NOS | negative |
| **TCGA-CV-5435** | male | stage iva | Larynx, NOS | negative |
| **TCGA-CV-5436** | male | stage iva | Floor of mouth, NOS | negative |
| **TCGA-CV-5439** | male | stage iva | Base of tongue, NOS | negative |
| **TCGA-CV-5440** | male | stage iva | Larynx, NOS | negative |
| **TCGA-CV-5441** | male | stage iva | Larynx, NOS | negative |
| **TCGA-CV-5442** | female | stage iva | Hard palate | positive |
| **TCGA-CV-5443** | male | stage iii | Larynx, NOS | positive |
| **TCGA-CV-5444** | male | stage iva | Larynx, NOS | negative |
| **TCGA-CV-5966** | female | stage iva | Overlapping lesion of lip, oral cavity and pharynx | negative |
| **TCGA-CV-5970** | male | stage iva | Tongue, NOS | positive |
| **TCGA-CV-5971** | male | stage iva | Tongue, NOS | positive |
| **TCGA-CV-5973** | female | stage iii | Tongue, NOS | negative |
| **TCGA-CV-5976** | male | stage iva | Tongue, NOS | negative |
| **TCGA-CV-5977** | male | stage iva | Tongue, NOS | negative |
| **TCGA-CV-5978** | female | stage ivb | Larynx, NOS | negative |
| **TCGA-CV-5979** | male | stage iva | Tongue, NOS | negative |
| **TCGA-CV-6003** | female | stage iii | Tongue, NOS | negative |
| **TCGA-CV-6433** | male | stage ii | Tongue, NOS | positive |
| **TCGA-CV-6436** | male | stage iva | Tongue, NOS | negative |
| **TCGA-CV-6441** | male | stage iii | Tongue, NOS | negative |
| **TCGA-CV-6933** | male | stage iii | Tongue, NOS | negative |
| **TCGA-CV-6934** | female | stage iva | Tongue, NOS | negative |

| | | | | |
|---|---|---|---|---|
| **TCGA-CV-6935** | male | stage iva | Larynx, NOS | negative |
| **TCGA-CV-6936** | male | stage iva | Floor of mouth, NOS | negative |
| **TCGA-CV-6937** | male | stage ii | Overlapping lesion of lip, oral cavity and pharynx | negative |
| **TCGA-CV-6938** | male | stage ii | Overlapping lesion of lip, oral cavity and pharynx | negative |
| **TCGA-CV-6939** | male | stage iva | Tongue, NOS | positive |
| **TCGA-CV-6940** | female | stage iii | Cheek mucosa | negative |
| **TCGA-CV-6941** | male | stage iii | Tongue, NOS | negative |
| **TCGA-CV-6942** | female | stage ii | Overlapping lesion of lip, oral cavity and pharynx | negative |
| **TCGA-CV-6943** | male | stage iii | Base of tongue, NOS | negative |
| **TCGA-CV-6945** | male | stage iva | Tongue, NOS | negative |
| **TCGA-CV-6948** | female | stage ivb | Floor of mouth, NOS | negative |
| **TCGA-CV-6950** | male | stage iva | Base of tongue, NOS | negative |
| **TCGA-CV-6951** | male | stage iva | Tongue, NOS | negative |
| **TCGA-CV-6952** | female | stage iva | Tongue, NOS | negative |
| **TCGA-CV-6953** | female | stage iii | Floor of mouth, NOS | negative |
| **TCGA-CV-6954** | male | stage iva | Tongue, NOS | negative |
| **TCGA-CV-6955** | female | stage ii | Overlapping lesion of lip, oral cavity and pharynx | negative |
| **TCGA-CV-6956** | male | stage iii | Tongue, NOS | negative |
| **TCGA-CV-6959** | male | stage iii | Tongue, NOS | negative |
| **TCGA-CV-6960** | male | stage iii | Overlapping lesion of lip, oral cavity and pharynx | negative |
| **TCGA-CV-6961** | male | stage ii | Tongue, NOS | positive |
| **TCGA-CV-6962** | male | stage iva | Larynx, NOS | negative |
| **TCGA-CV-7089** | male | stage iva | Larynx, NOS | negative |
| **TCGA-CV-7090** | male | stage ii | Overlapping lesion of lip, oral cavity and pharynx | negative |
| **TCGA-CV-7091** | male | stage i | Overlapping lesion of lip, oral cavity and pharynx | negative |
| **TCGA-CV-7095** | female | stage ii | Overlapping lesion of lip, oral cavity and pharynx | negative |
| **TCGA-CV-7097** | male | stage ii | Overlapping lesion of lip, oral cavity and pharynx | negative |

| | | | | |
|---|---|---|---|---|
| **TCGA-CV-7099** | female | stage ii | Overlapping lesion of lip, oral cavity and pharynx | negative |
| **TCGA-CV-7100** | male | stage iii | Overlapping lesion of lip, oral cavity and pharynx | positive |
| **TCGA-CV-7101** | male | stage ii | Larynx, NOS | negative |
| **TCGA-CV-7102** | female | stage iva | Floor of mouth, NOS | negative |
| **TCGA-CV-7103** | male | stage iva | Tongue, NOS | negative |
| **TCGA-CV-7104** | female | stage iva | Tongue, NOS | negative |
| **TCGA-CV-7177** | female | stage ii | Larynx, NOS | negative |
| **TCGA-CV-7178** | female | stage iva | Overlapping lesion of lip, oral cavity and pharynx | negative |
| **TCGA-CV-7180** | male | stage ii | Tongue, NOS | negative |
| **TCGA-CV-7183** | male | stage ii | Overlapping lesion of lip, oral cavity and pharynx | negative |
| **TCGA-CV-7235** | male | stage ii | Floor of mouth, NOS | negative |
| **TCGA-CV-7236** | female | stage iva | Tongue, NOS | negative |
| **TCGA-CV-7238** | female | stage ii | Tongue, NOS | negative |
| **TCGA-CV-7242** | female | stage ii | Larynx, NOS | negative |
| **TCGA-CV-7243** | male | stage iii | Tongue, NOS | negative |
| **TCGA-CV-7245** | male | stage iva | Larynx, NOS | negative |
| **TCGA-CV-7247** | male | stage iii | Larynx, NOS | negative |
| **TCGA-CV-7248** | female | stage iva | Larynx, NOS | negative |
| **TCGA-CV-7250** | male | stage iva | Larynx, NOS | negative |
| **TCGA-CV-7252** | female | stage iva | Overlapping lesion of lip, oral cavity and pharynx | negative |
| **TCGA-CV-7253** | male | stage iva | Overlapping lesion of lip, oral cavity and pharynx | negative |
| **TCGA-CV-7254** | male | stage ii | Overlapping lesion of lip, oral cavity and pharynx | negative |
| **TCGA-CV-7255** | female | stage iva | Tongue, NOS | negative |
| **TCGA-CV-7261** | male | stage iva | Larynx, NOS | negative |
| **TCGA-CV-7263** | male | stage ii | Overlapping lesion of lip, oral cavity and pharynx | negative |
| **TCGA-CV-7406** | male | stage ii | Base of tongue, NOS | positive |
| **TCGA-CV-7407** | female | stage ii | Floor of mouth, NOS | negative |

| | | | | |
|---|---|---|---|---|
| **TCGA-CV-7409** | male | stage ivb | Overlapping lesion of lip, oral cavity and pharynx | negative |
| **TCGA-CV-7410** | male | stage iii | Larynx, NOS | negative |
| **TCGA-CV-7411** | female | stage iva | Overlapping lesion of lip, oral cavity and pharynx | negative |
| **TCGA-CV-7413** | female | stage ii | Overlapping lesion of lip, oral cavity and pharynx | negative |
| **TCGA-CV-7414** | male | stage iva | Overlapping lesion of lip, oral cavity and pharynx | negative |
| **TCGA-CV-7415** | male | stage iva | Larynx, NOS | negative |
| **TCGA-CV-7416** | female | stage iva | Overlapping lesion of lip, oral cavity and pharynx | negative |
| **TCGA-CV-7418** | male | stage iva | Larynx, NOS | negative |
| **TCGA-CV-7421** | male | stage iva | Larynx, NOS | negative |
| **TCGA-CV-7422** | female | stage iva | Larynx, NOS | negative |
| **TCGA-CV-7423** | male | stage ii | Overlapping lesion of lip, oral cavity and pharynx | negative |
| **TCGA-CV-7424** | male | stage iva | Larynx, NOS | negative |
| **TCGA-CV-7425** | female | stage iii | Overlapping lesion of lip, oral cavity and pharynx | negative |
| **TCGA-CV-7427** | female | stage ii | Overlapping lesion of lip, oral cavity and pharynx | negative |
| **TCGA-CV-7428** | male | stage iva | Overlapping lesion of lip, oral cavity and pharynx | negative |
| **TCGA-CV-7429** | male | stage iva | Overlapping lesion of lip, oral cavity and pharynx | negative |
| **TCGA-CV-7430** | male | stage iva | Larynx, NOS | negative |
| **TCGA-CV-7432** | male | stage ii | Overlapping lesion of lip, oral cavity and pharynx | negative |
| **TCGA-CV-7433** | male | stage iva | Larynx, NOS | negative |
| **TCGA-CV-7434** | male | stage iva | Overlapping lesion of lip, oral cavity and pharynx | negative |
| **TCGA-CV-7435** | female | stage iva | Overlapping lesion of lip, oral cavity and pharynx | negative |
| **TCGA-CV-7437** | male | stage ii | Larynx, NOS | negative |
| **TCGA-CV-7438** | female | stage i | Tongue, NOS | negative |
| **TCGA-CV-7440** | male | stage ii | Larynx, NOS | negative |
| **TCGA-CV-7446** | male | stage iva | Tongue, NOS | negative |
| **TCGA-CV-7568** | female | stage iva | Overlapping lesion of lip, oral cavity and pharynx | negative |
| **TCGA-CV-A45O** | male | N/A | Gum, NOS | negative |

| | | | | |
|---|---|---|---|---|
| **TCGA-CV-A45P** | female | stage i | Tongue, NOS | negative |
| **TCGA-CV-A45Q** | female | N/A | Mouth, NOS | negative |
| **TCGA-CV-A45R** | male | stage iii | Tongue, NOS | negative |
| **TCGA-CV-A45T** | female | N/A | Tongue, NOS | negative |
| **TCGA-CV-A45U** | male | stage iva | Mouth, NOS | negative |
| **TCGA-CV-A45V** | female | stage iva | Mouth, NOS | negative |
| **TCGA-CV-A45W** | male | stage ii | Larynx, NOS | negative |
| **TCGA-CV-A45X** | male | stage iva | Floor of mouth, NOS | negative |
| **TCGA-CV-A45Y** | male | stage iva | Larynx, NOS | negative |
| **TCGA-CV-A45Z** | male | N/A | Larynx, NOS | negative |
| **TCGA-CV-A460** | male | N/A | Larynx, NOS | negative |
| **TCGA-CV-A461** | male | N/A | Larynx, NOS | negative |
| **TCGA-CV-A463** | female | stage iva | Floor of mouth, NOS | negative |
| **TCGA-CV-A464** | male | stage iii | Cheek mucosa | negative |
| **TCGA-CV-A465** | male | stage iii | Tongue, NOS | negative |
| **TCGA-CV-A468** | male | stage iva | Lip, NOS | negative |
| **TCGA-CV-A6JD** | female | N/A | Floor of mouth, NOS | negative |
| **TCGA-CV-A6JE** | male | N/A | Mouth, NOS | negative |
| **TCGA-CV-A6JM** | male | N/A | Pharynx, NOS | negative |
| **TCGA-CV-A6JN** | male | N/A | Mouth, NOS | negative |
| **TCGA-CV-A6JO** | male | stage iva | Tongue, NOS | negative |
| **TCGA-CV-A6JT** | male | stage ii | Tongue, NOS | negative |
| **TCGA-CV-A6JU** | female | stage ivb | Tongue, NOS | negative |
| **TCGA-CV-A6JY** | male | stage iva | Mouth, NOS | negative |
| **TCGA-CV-A6JZ** | male | stage ii | Mouth, NOS | negative |
| **TCGA-CV-A6K0** | male | stage i | Tongue, NOS | negative |
| **TCGA-CV-A6K1** | male | stage iva | Larynx, NOS | negative |

| | | | | |
|---|---|---|---|---|
| **TCGA-CV-A6K2** | male | stage iva | Mouth, NOS | negative |
| **TCGA-CX-7086** | male | stage iii | Floor of mouth, NOS | negative |
| **TCGA-CX-7219** | male | stage iva | Floor of mouth, NOS | negative |
| **TCGA-CX-A4AQ** | male | stage iva | Floor of mouth, NOS | negative |
| **TCGA-D6-6515** | female | stage ii | Tongue, NOS | negative |
| **TCGA-D6-6516** | male | stage i | Lip, NOS | negative |
| **TCGA-D6-6517** | male | stage iii | Larynx, NOS | negative |
| **TCGA-D6-6823** | male | stage ii | Tongue, NOS | negative |
| **TCGA-D6-6824** | male | stage iva | Larynx, NOS | negative |
| **TCGA-D6-6825** | male | stage i | Tongue, NOS | negative |
| **TCGA-D6-6826** | female | stage iva | Larynx, NOS | negative |
| **TCGA-D6-6827** | female | stage i | Lip, NOS | negative |
| **TCGA-D6-8568** | male | stage ii | Larynx, NOS | negative |
| **TCGA-D6-8569** | male | stage ii | Tongue, NOS | negative |
| **TCGA-D6-A4Z9** | male | stage iva | Tongue, NOS | negative |
| **TCGA-D6-A4ZB** | male | stage iii | Tongue, NOS | negative |
| **TCGA-D6-A6EN** | male | stage iii | Cheek mucosa | negative |
| **TCGA-D6-A6EO** | male | stage iva | Floor of mouth, NOS | negative |
| **TCGA-D6-A6EP** | male | stage iii | Hypopharynx, NOS | negative |
| **TCGA-D6-A6EQ** | male | stage iva | Larynx, NOS | negative |
| **TCGA-D6-A74Q** | male | stage iva | Larynx, NOS | negative |
| **TCGA-DQ-5624** | female | N/A | Tongue, NOS | negative |
| **TCGA-DQ-5625** | female | N/A | Tongue, NOS | negative |
| **TCGA-DQ-5629** | male | N/A | Larynx, NOS | negative |
| **TCGA-DQ-5630** | male | N/A | Tongue, NOS | negative |
| **TCGA-DQ-5631** | male | N/A | Tongue, NOS | negative |
| **TCGA-DQ-7588** | male | N/A | Overlapping lesion of lip, oral cavity and pharynx | negative |

| | | | | |
|---|---|---|---|---|
| **TCGA-DQ-7589** | male | N/A | Larynx, NOS | positive |
| **TCGA-DQ-7590** | male | N/A | Tonsil, NOS | positive |
| **TCGA-DQ-7591** | male | N/A | Base of tongue, NOS | positive |
| **TCGA-DQ-7592** | male | N/A | Tongue, NOS | negative |
| **TCGA-DQ-7593** | male | N/A | Base of tongue, NOS | positive |
| **TCGA-DQ-7594** | male | N/A | Base of tongue, NOS | positive |
| **TCGA-DQ-7595** | male | N/A | Larynx, NOS | negative |
| **TCGA-DQ-7596** | male | N/A | Tonsil, NOS | positive |
| **TCGA-F7-7848** | male | stage iva | Larynx, NOS | negative |
| **TCGA-F7-8298** | male | stage i | Larynx, NOS | negative |
| **TCGA-F7-8489** | male | stage ii | Floor of mouth, NOS | negative |
| **TCGA-F7-A50G** | male | stage iii | Tongue, NOS | negative |
| **TCGA-F7-A50I** | male | stage iva | Larynx, NOS | negative |
| **TCGA-F7-A50J** | female | stage iii | Tongue, NOS | negative |
| **TCGA-F7-A61S** | male | stage iii | Tongue, NOS | negative |
| **TCGA-F7-A61V** | male | stage ii | Base of tongue, NOS | negative |
| **TCGA-F7-A61W** | male | stage iva | Tongue, NOS | negative |
| **TCGA-F7-A620** | male | stage iii | Base of tongue, NOS | negative |
| **TCGA-F7-A622** | male | stage iii | Larynx, NOS | negative |
| **TCGA-F7-A623** | male | stage iva | Larynx, NOS | negative |
| **TCGA-F7-A624** | male | stage ii | Cheek mucosa | negative |
| **TCGA-H7-A6C4** | female | stage iva | Tongue, NOS | negative |
| **TCGA-H7-A76A** | male | N/A | Tonsil, NOS | positive |
| **TCGA-HD-7229** | male | stage iva | Larynx, NOS | negative |
| **TCGA-HD-7753** | male | stage i | Oropharynx, NOS | negative |
| **TCGA-HD-7754** | male | stage iva | Tonsil, NOS | positive |
| **TCGA-HD-7831** | male | stage iva | Tongue, NOS | negative |

| | | | | |
|---|---|---|---|---|
| **TCGA-HD-7832** | male | stage iva | Floor of mouth, NOS | positive |
| **TCGA-HD-7917** | male | stage ii | Floor of mouth, NOS | negative |
| **TCGA-HD-8224** | male | stage iva | Base of tongue, NOS | negative |
| **TCGA-HD-8314** | male | stage iii | Base of tongue, NOS | positive |
| **TCGA-HD-8634** | female | stage i | Tongue, NOS | negative |
| **TCGA-HD-A4C1** | female | stage iva | Cheek mucosa | negative |
| **TCGA-HD-A633** | male | stage iva | Mouth, NOS | positive |
| **TCGA-HD-A634** | male | stage iva | Tonsil, NOS | positive |
| **TCGA-HD-A6HZ** | female | stage iii | Tongue, NOS | negative |
| **TCGA-HD-A6I0** | male | stage iva | Mouth, NOS | negative |
| **TCGA-HL-7533** | male | N/A | Overlapping lesion of lip, oral cavity and pharynx | positive |
| **TCGA-IQ-7630** | male | stage ii | Oropharynx, NOS | negative |
| **TCGA-IQ-A61E** | female | stage iii | Tongue, NOS | negative |
| **TCGA-IQ-A61G** | male | stage iva | Floor of mouth, NOS | negative |
| **TCGA-IQ-A61H** | male | stage ii | Tongue, NOS | negative |
| **TCGA-IQ-A61I** | male | stage iva | Tonsil, NOS | positive |
| **TCGA-IQ-A61J** | male | stage iva | Tongue, NOS | negative |
| **TCGA-IQ-A61O** | male | stage iva | Oropharynx, NOS | negative |
| **TCGA-IQ-A6SG** | female | stage iii | Tongue, NOS | negative |
| **TCGA-IQ-A6SH** | male | stage iii | Tongue, NOS | negative |
| **TCGA-KU-A66S** | female | stage iii | Larynx, NOS | negative |
| **TCGA-KU-A66T** | female | stage iva | Floor of mouth, NOS | negative |
| **TCGA-KU-A6H7** | female | stage iva | Tonsil, NOS | positive |
| **TCGA-KU-A6H8** | male | stage iva | Tongue, NOS | negative |
| **TCGA-MT-A51W** | female | stage i | Tonsil, NOS | negative |
| **TCGA-MT-A51X** | male | stage iva | Tongue, NOS | negative |
| **TCGA-MT-A67A** | female | stage i | Tongue, NOS | negative |

| | | | | |
|---|---|---|---|---|
| **TCGA-MT-A67D** | male | stage ii | Anterior floor of mouth | negative |
| **TCGA-MT-A67F** | female | stage iva | Mouth, NOS | negative |
| **TCGA-MT-A7BN** | male | stage iva | Floor of mouth, NOS | negative |
| **TCGA-MZ-A6I9** | male | N/A | Oropharynx, NOS | positive |
| **TCGA-MZ-A7D7** | male | stage iva | Base of tongue, NOS | negative |
| **TCGA-P3-A5Q5** | male | stage iva | Tonsil, NOS | positive |
| **TCGA-P3-A5Q6** | male | stage iva | Tonsil, NOS | negative |
| **TCGA-P3-A5QA** | male | stage i | Tongue, NOS | negative |
| **TCGA-P3-A5QE** | male | stage iii | Base of tongue, NOS | positive |
| **TCGA-P3-A5QF** | male | stage iva | Upper Gum | positive |
| **TCGA-P3-A6SW** | male | stage ivb | Tonsil, NOS | positive |
| **TCGA-P3-A6SX** | male | stage iii | Tonsil, NOS | negative |
| **TCGA-P3-A6T0** | female | stage iva | Floor of mouth, NOS | negative |
| **TCGA-P3-A6T3** | male | stage iva | Mouth, NOS | negative |
| **TCGA-P3-A6T4** | male | stage iva | Floor of mouth, NOS | negative |
| **TCGA-P3-A6T5** | female | stage iva | Lower gum | negative |
| **TCGA-QK-A64Z** | female | stage iva | Hard palate | negative |
| **TCGA-QK-A652** | male | stage iii | Tongue, NOS | negative |
| **TCGA-QK-A6IF** | male | N/A | Tonsil, NOS | positive |
| **TCGA-QK-A6IG** | male | stage iii | Cheek mucosa | negative |
| **TCGA-QK-A6IH** | female | stage ivb | Gum, NOS | negative |
| **TCGA-QK-A6II** | male | stage iva | Floor of mouth, NOS | negative |
| **TCGA-QK-A6IJ** | male | stage iii | Floor of mouth, NOS | negative |
| **TCGA-QK-A6V9** | male | stage ii | Tonsil, NOS | positive |
| **TCGA-QK-A6VB** | male | stage iva | Floor of mouth, NOS | negative |
| **TCGA-QK-A6VC** | female | stage iva | Hypopharynx, NOS | negative |
| **TCGA-QK-A8Z7** | male | stage iva | Floor of mouth, NOS | negative |

| | | | | |
|---|---|---|---|---|
| **TCGA-QK-A8Z8** | female | stage ivc | Larynx, NOS | negative |
| **TCGA-QK-A8Z9** | male | stage iva | Floor of mouth, NOS | negative |
| **TCGA-QK-A8ZA** | male | N/A | Oropharynx, NOS | negative |
| **TCGA-QK-A8ZB** | male | stage iva | Larynx, NOS | negative |
| **TCGA-QK-AA3J** | male | stage i | Larynx, NOS | negative |
| **TCGA-RS-A6TO** | female | stage iva | Mouth, NOS | negative |
| **TCGA-RS-A6TP** | male | stage ii | Tonsil, NOS | positive |
| **TCGA-T2-A6WX** | female | N/A | Floor of mouth, NOS | negative |
| **TCGA-T2-A6WZ** | male | N/A | Base of tongue, NOS | negative |
| **TCGA-T2-A6X0** | male | N/A | Tonsil, NOS | positive |
| **TCGA-T2-A6X2** | male | stage iii | Gum, NOS | negative |
| **TCGA-T3-A92M** | male | stage iva | Larynx, NOS | negative |
| **TCGA-TN-A7HI** | male | stage iva | Tonsil, NOS | positive |
| **TCGA-TN-A7HJ** | male | stage iva | Larynx, NOS | negative |
| **TCGA-TN-A7HL** | male | stage iva | Hypopharynx, NOS | positive |
| **TCGA-UF-A718** | male | stage iva | Larynx, NOS | negative |
| **TCGA-UF-A719** | male | stage ii | Floor of mouth, NOS | negative |
| **TCGA-UF-A71A** | male | stage iva | Floor of mouth, NOS | negative |
| **TCGA-UF-A71B** | male | stage iva | Gum, NOS | negative |
| **TCGA-UF-A71D** | female | stage iva | Larynx, NOS | negative |
| **TCGA-UF-A71E** | male | stage iva | Floor of mouth, NOS | negative |
| **TCGA-UF-A7J9** | male | stage iva | Larynx, NOS | negative |
| **TCGA-UF-A7JA** | female | stage iva | Cheek mucosa | negative |
| **TCGA-UF-A7JC** | male | stage iva | Floor of mouth, NOS | negative |
| **TCGA-UF-A7JD** | male | stage iva | Cheek mucosa | negative |
| **TCGA-UF-A7JF** | male | stage iva | Larynx, NOS | negative |
| **TCGA-UF-A7JH** | male | stage iva | Larynx, NOS | negative |

| | | | | |
|---|---|---|---|---|
| **TCGA-UF-A7JJ** | male | stage iva | Larynx, NOS | negative |
| **TCGA-UF-A7JK** | male | stage iva | Larynx, NOS | negative |
| **TCGA-UF-A7JO** | female | stage iva | Floor of mouth, NOS | negative |
| **TCGA-UF-A7JS** | male | stage iva | Tongue, NOS | negative |
| **TCGA-UF-A7JT** | female | stage iva | Floor of mouth, NOS | negative |
| **TCGA-UF-A7JV** | female | stage iva | Hypopharynx, NOS | negative |
| **TCGA-WA-A7GZ** | male | N/A | Floor of mouth, NOS | negative |
| **TCGA-WA-A7H4** | male | stage ii | Tongue, NOS | negative |

**Suppl. Table 2: Clinical characterization of TCGA-HNSC (N=412 patients)** NOS = not otherwise specified

| Patient | Sex | Stage | Site of origin | isEBV |
|---|---|---|---|---|
| **TCGA-3M-AB46** | male | stage ib | Lesser curvature of stomach, NOS | nonEBV |
| **TCGA-3M-AB47** | male | stage iiib | Stomach, NOS | nonEBV |
| **TCGA-B7-5816** | female | stage iib | Body of stomach | nonEBV |
| **TCGA-BR-6452** | female | stage iia | Body of stomach | nonEBV |
| **TCGA-BR-6453** | male | stage iia | Body of stomach | nonEBV |
| **TCGA-BR-6454** | male | stage iia | Cardia, NOS | nonEBV |
| **TCGA-BR-6455** | male | stage iib | Body of stomach | EBV |
| **TCGA-BR-6456** | female | stage iib | Body of stomach | nonEBV |
| **TCGA-BR-6457** | male | stage iia | Body of stomach | nonEBV |
| **TCGA-BR-6458** | female | stage iib | Body of stomach | nonEBV |
| **TCGA-BR-6563** | male | stage iib | Gastric antrum | nonEBV |
| **TCGA-BR-6564** | female | stage iiia | Body of stomach | nonEBV |
| **TCGA-BR-6565** | male | stage iib | Body of stomach | nonEBV |
| **TCGA-BR-6566** | female | stage iia | Body of stomach | nonEBV |
| **TCGA-BR-6705** | female | stage iiib | Body of stomach | nonEBV |
| **TCGA-BR-6706** | male | stage iiia | Cardia, NOS | EBV |
| **TCGA-BR-6707** | male | stage iia | Body of stomach | EBV |
| **TCGA-BR-6801** | male | stage iia | Body of stomach | nonEBV |
| **TCGA-BR-6802** | male | stage iiia | Body of stomach | nonEBV |
| **TCGA-BR-6803** | female | stage iia | Body of stomach | nonEBV |
| **TCGA-BR-6852** | female | stage iia | Cardia, NOS | nonEBV |
| **TCGA-BR-7196** | male | stage iv | Cardia, NOS | EBV |
| **TCGA-BR-7197** | male | stage ii | Gastric antrum | nonEBV |
| **TCGA-BR-7703** | male | stage ia | Body of stomach | nonEBV |
| **TCGA-BR-7707** | female | stage ib | Body of stomach | nonEBV |
| **TCGA-BR-7715** | male | stage iia | Body of stomach | nonEBV |

| | | | | |
|---|---|---|---|---|
| **TCGA-BR-7716** | female | stage iib | Body of stomach | nonEBV |
| **TCGA-BR-7717** | male | stage iv | Cardia, NOS | nonEBV |
| **TCGA-BR-7722** | male | stage iib | Cardia, NOS | nonEBV |
| **TCGA-BR-7723** | male | stage iiib | Body of stomach | nonEBV |
| **TCGA-BR-7851** | male | stage iib | Body of stomach | nonEBV |
| **TCGA-BR-7901** | male | stage iib | Cardia, NOS | nonEBV |
| **TCGA-BR-7957** | female | stage iv | Body of stomach | nonEBV |
| **TCGA-BR-7958** | male | stage iiib | Body of stomach | EBV |
| **TCGA-BR-7959** | male | stage iiia | Cardia, NOS | nonEBV |
| **TCGA-BR-8058** | female | stage iiib | Gastric antrum | nonEBV |
| **TCGA-BR-8059** | male | stage iii | Fundus of stomach | nonEBV |
| **TCGA-BR-8077** | female | stage iiib | Body of stomach | nonEBV |
| **TCGA-BR-8080** | female | stage iiic | Body of stomach | nonEBV |
| **TCGA-BR-8081** | female | stage iib | Body of stomach | nonEBV |
| **TCGA-BR-8284** | female | stage iiic | Gastric antrum | nonEBV |
| **TCGA-BR-8286** | male | stage ii | Gastric antrum | nonEBV |
| **TCGA-BR-8289** | male | stage iv | Body of stomach | nonEBV |
| **TCGA-BR-8291** | male | stage iib | Gastric antrum | nonEBV |
| **TCGA-BR-8297** | male | stage iiic | Body of stomach | nonEBV |
| **TCGA-BR-8361** | female | stage iiic | Fundus of stomach | nonEBV |
| **TCGA-BR-8363** | female | stage ib | Gastric antrum | nonEBV |
| **TCGA-BR-8364** | female | stage iiic | Fundus of stomach | nonEBV |
| **TCGA-BR-8365** | female | stage iia | Fundus of stomach | nonEBV |
| **TCGA-BR-8366** | female | stage iia | Fundus of stomach | EBV |
| **TCGA-BR-8367** | male | stage iiib | Fundus of stomach | nonEBV |
| **TCGA-BR-8368** | female | stage ib | Fundus of stomach | nonEBV |
| **TCGA-BR-8369** | female | stage iiib | Gastric antrum | nonEBV |

| | | | | |
|---|---|---|---|---|
| **TCGA-BR-8371** | male | stage iiib | Cardia, NOS | nonEBV |
| **TCGA-BR-8372** | male | stage iiic | Fundus of stomach | nonEBV |
| **TCGA-BR-8373** | female | stage iiia | Fundus of stomach | nonEBV |
| **TCGA-BR-8380** | male | stage iiic | Gastric antrum | nonEBV |
| **TCGA-BR-8381** | male | stage iib | Fundus of stomach | EBV |
| **TCGA-BR-8382** | female | stage iiic | Gastric antrum | nonEBV |
| **TCGA-BR-8384** | male | stage iiic | Gastric antrum | nonEBV |
| **TCGA-BR-8483** | male | stage iiia | Gastric antrum | nonEBV |
| **TCGA-BR-8484** | male | stage iiia | Fundus of stomach | nonEBV |
| **TCGA-BR-8485** | female | stage iiic | Fundus of stomach | nonEBV |
| **TCGA-BR-8486** | female | stage ia | Fundus of stomach | nonEBV |
| **TCGA-BR-8487** | female | stage iia | Gastric antrum | nonEBV |
| **TCGA-BR-8588** | female | stage iib | Fundus of stomach | nonEBV |
| **TCGA-BR-8589** | male | stage iiib | Fundus of stomach | EBV |
| **TCGA-BR-8590** | male | stage iiic | Fundus of stomach | nonEBV |
| **TCGA-BR-8591** | male | stage iiic | Gastric antrum | nonEBV |
| **TCGA-BR-8592** | female | stage iiib | Fundus of stomach | nonEBV |
| **TCGA-BR-8677** | female | stage iiib | Gastric antrum | nonEBV |
| **TCGA-BR-8678** | male | stage ib | Fundus of stomach | nonEBV |
| **TCGA-BR-8679** | female | stage ib | Fundus of stomach | nonEBV |
| **TCGA-BR-8680** | male | stage iv | Gastric antrum | nonEBV |
| **TCGA-BR-8682** | male | stage iib | Gastric antrum | nonEBV |
| **TCGA-BR-8683** | male | stage iiib | Gastric antrum | nonEBV |
| **TCGA-BR-8686** | male | stage iiib | Gastric antrum | EBV |
| **TCGA-BR-8687** | female | stage iiic | Cardia, NOS | nonEBV |
| **TCGA-BR-A44T** | female | stage iia | Cardia, NOS | nonEBV |
| **TCGA-BR-A44U** | male | stage iiib | Gastric antrum | nonEBV |

| | | | | |
|---|---|---|---|---|
| **TCGA-BR-A452** | male | stage iiia | Gastric antrum | nonEBV |
| **TCGA-BR-A453** | male | stage iv | Gastric antrum | nonEBV |
| **TCGA-BR-A4CR** | female | stage iiic | Fundus of stomach | nonEBV |
| **TCGA-BR-A4CS** | male | stage iiic | Fundus of stomach | nonEBV |
| **TCGA-BR-A4IU** | female | stage iiia | Gastric antrum | nonEBV |
| **TCGA-BR-A4IV** | male | stage iiib | Gastric antrum | nonEBV |
| **TCGA-BR-A4IY** | male | stage iib | Gastric antrum | nonEBV |
| **TCGA-BR-A4IZ** | female | stage iiib | Gastric antrum | nonEBV |
| **TCGA-BR-A4J1** | male | stage iiia | Gastric antrum | nonEBV |
| **TCGA-BR-A4J2** | male | stage iib | Gastric antrum | nonEBV |
| **TCGA-BR-A4J4** | male | stage iiib | Gastric antrum | EBV |
| **TCGA-BR-A4J5** | male | stage iiia | Gastric antrum | nonEBV |
| **TCGA-BR-A4J6** | female | stage iia | Cardia, NOS | nonEBV |
| **TCGA-BR-A4J7** | male | stage iib | Gastric antrum | nonEBV |
| **TCGA-BR-A4J8** | female | stage iiib | Cardia, NOS | nonEBV |
| **TCGA-BR-A4J9** | male | stage iia | Gastric antrum | nonEBV |
| **TCGA-BR-A4PD** | female | stage iib | Fundus of stomach | nonEBV |
| **TCGA-BR-A4PE** | female | stage ib | Cardia, NOS | nonEBV |
| **TCGA-BR-A4PF** | male | stage iiib | Gastric antrum | nonEBV |
| **TCGA-BR-A4QI** | female | stage iia | Gastric antrum | nonEBV |
| **TCGA-CD-5798** | male | stage ii | Gastric antrum | nonEBV |
| **TCGA-CD-5799** | male | stage ii | Gastric antrum | nonEBV |
| **TCGA-CD-5800** | female | stage ii | Gastric antrum | nonEBV |
| **TCGA-CD-5801** | male | stage iiia | Gastric antrum | EBV |
| **TCGA-CD-5803** | female | stage ii | Gastric antrum | nonEBV |
| **TCGA-CD-5804** | male | not reported | Pylorus | nonEBV |
| **TCGA-CD-8524** | female | stage ii | Gastric antrum | nonEBV |

| | | | | |
|---|---|---|---|---|
| **TCGA-CD-8526** | female | stage iiia | Fundus of stomach | nonEBV |
| **TCGA-CD-8528** | female | stage iiia | Gastric antrum | nonEBV |
| **TCGA-CD-8529** | male | stage iv | Gastric antrum | nonEBV |
| **TCGA-CD-8530** | male | stage ii | Fundus of stomach | nonEBV |
| **TCGA-CD-8531** | female | stage iiia | Fundus of stomach | nonEBV |
| **TCGA-CD-8532** | male | stage ii | Cardia, NOS | nonEBV |
| **TCGA-CD-8534** | male | stage ii | Gastric antrum | nonEBV |
| **TCGA-CD-8535** | male | stage iiia | Fundus of stomach | nonEBV |
| **TCGA-CD-8536** | male | stage ii | Fundus of stomach | nonEBV |
| **TCGA-CD-A486** | male | stage iia | Cardia, NOS | nonEBV |
| **TCGA-CD-A487** | male | stage iib | Gastric antrum | nonEBV |
| **TCGA-CD-A489** | male | stage iia | Gastric antrum | nonEBV |
| **TCGA-CD-A48A** | male | stage iia | Cardia, NOS | nonEBV |
| **TCGA-CD-A48C** | female | stage iib | Gastric antrum | nonEBV |
| **TCGA-CD-A4MG** | male | stage iia | Cardia, NOS | nonEBV |
| **TCGA-CD-A4MH** | female | stage iia | Gastric antrum | nonEBV |
| **TCGA-CD-A4MI** | male | stage iiia | Gastric antrum | nonEBV |
| **TCGA-CD-A4MJ** | male | stage ib | Fundus of stomach | nonEBV |
| **TCGA-CG-4304** | male | stage ib | Cardia, NOS | nonEBV |
| **TCGA-CG-4305** | male | stage ii | Gastric antrum | nonEBV |
| **TCGA-CG-4306** | male | stage iv | Gastric antrum | nonEBV |
| **TCGA-CG-4438** | male | stage iv | Gastric antrum | nonEBV |
| **TCGA-CG-4440** | female | stage iv | Body of stomach | nonEBV |
| **TCGA-CG-4441** | male | stage iiia | Body of stomach | nonEBV |
| **TCGA-CG-4444** | male | stage iiia | Body of stomach | nonEBV |
| **TCGA-CG-4449** | male | stage ii | Gastric antrum | nonEBV |
| **TCGA-CG-4465** | female | stage iv | Gastric antrum | nonEBV |

| | | | | |
|---|---|---|---|---|
| **TCGA-CG-4466** | female | stage ib | Cardia, NOS | nonEBV |
| **TCGA-CG-4469** | male | stage iv | Stomach, NOS | nonEBV |
| **TCGA-CG-4474** | female | stage iv | Gastric antrum | nonEBV |
| **TCGA-CG-4475** | male | stage iib | Stomach, NOS | nonEBV |
| **TCGA-CG-4476** | male | stage iiic | Body of stomach | nonEBV |
| **TCGA-CG-4477** | female | stage ib | Cardia, NOS | nonEBV |
| **TCGA-D7-5577** | female | stage iiia | Cardia, NOS | EBV |
| **TCGA-D7-5578** | male | stage iiia | Body of stomach | nonEBV |
| **TCGA-D7-6518** | male | stage iiia | Cardia, NOS | nonEBV |
| **TCGA-D7-6519** | female | not reported | Cardia, NOS | nonEBV |
| **TCGA-D7-6520** | male | stage iiia | Gastric antrum | nonEBV |
| **TCGA-D7-6521** | male | not reported | Body of stomach | nonEBV |
| **TCGA-D7-6522** | male | stage ib | Gastric antrum | nonEBV |
| **TCGA-D7-6524** | male | stage ii | Gastric antrum | nonEBV |
| **TCGA-D7-6525** | male | stage iiia | Cardia, NOS | nonEBV |
| **TCGA-D7-6526** | female | stage iiia | Body of stomach | nonEBV |
| **TCGA-D7-6527** | male | stage ii | Cardia, NOS | nonEBV |
| **TCGA-D7-6528** | female | stage ib | Body of stomach | nonEBV |
| **TCGA-D7-6815** | female | stage iib | Body of stomach | nonEBV |
| **TCGA-D7-6817** | male | stage iiia | Body of stomach | nonEBV |
| **TCGA-D7-6818** | male | stage iiia | Body of stomach | nonEBV |
| **TCGA-D7-6820** | male | stage iib | Body of stomach | nonEBV |
| **TCGA-D7-6822** | male | stage ib | Body of stomach | nonEBV |
| **TCGA-D7-8570** | male | stage iiib | Cardia, NOS | EBV |
| **TCGA-D7-8572** | male | stage iib | Fundus of stomach | nonEBV |
| **TCGA-D7-8573** | male | stage iia | Fundus of stomach | EBV |
| **TCGA-D7-8574** | male | stage iiia | Gastric antrum | nonEBV |

| | | | | |
|---|---|---|---|---|
| **TCGA-D7-8575** | male | stage iiia | Fundus of stomach | nonEBV |
| **TCGA-D7-8576** | female | stage iiib | Gastric antrum | nonEBV |
| **TCGA-D7-8578** | male | stage ib | Gastric antrum | nonEBV |
| **TCGA-D7-8579** | female | stage iib | Gastric antrum | nonEBV |
| **TCGA-D7-A4YT** | male | stage iiia | Cardia, NOS | nonEBV |
| **TCGA-D7-A4YU** | male | stage iiib | Fundus of stomach | nonEBV |
| **TCGA-D7-A4YV** | female | stage iib | Gastric antrum | nonEBV |
| **TCGA-D7-A4YX** | male | stage iib | Fundus of stomach | EBV |
| **TCGA-D7-A4YY** | male | stage iiib | Gastric antrum | nonEBV |
| **TCGA-D7-A4Z0** | female | stage iib | Gastric antrum | nonEBV |
| **TCGA-D7-A6EV** | female | stage iib | Gastric antrum | nonEBV |
| **TCGA-D7-A6EX** | female | stage iiia | Gastric antrum | nonEBV |
| **TCGA-D7-A6EY** | female | stage iiib | Gastric antrum | nonEBV |
| **TCGA-D7-A6EZ** | male | stage iiia | Fundus of stomach | EBV |
| **TCGA-D7-A6F0** | female | stage ib | Cardia, NOS | nonEBV |
| **TCGA-D7-A747** | male | stage iib | Gastric antrum | nonEBV |
| **TCGA-D7-A748** | female | stage iv | Fundus of stomach | nonEBV |
| **TCGA-D7-A74A** | female | stage iiia | Cardia, NOS | nonEBV |
| **TCGA-EQ-A4SO** | male | stage iiib | Gastric antrum | nonEBV |
| **TCGA-F1-6177** | male | stage i | Gastric antrum | nonEBV |
| **TCGA-F1-6874** | male | stage ib | Cardia, NOS | nonEBV |
| **TCGA-F1-6875** | male | stage ia | Gastric antrum | nonEBV |
| **TCGA-F1-A448** | male | stage iiib | Gastric antrum | nonEBV |
| **TCGA-F1-A72C** | male | stage iia | Stomach, NOS | nonEBV |
| **TCGA-FP-7735** | male | stage ib | Cardia, NOS | nonEBV |
| **TCGA-FP-7829** | male | stage iib | Cardia, NOS | nonEBV |
| **TCGA-FP-7916** | male | stage iiic | Gastric antrum | EBV |

| | | | | |
|---|---|---|---|---|
| **TCGA-FP-7998** | male | stage iiic | Body of stomach | EBV |
| **TCGA-FP-8099** | male | stage iia | Cardia, NOS | nonEBV |
| **TCGA-FP-8209** | male | stage ib | Gastric antrum | nonEBV |
| **TCGA-FP-8210** | male | stage iiia | Gastric antrum | nonEBV |
| **TCGA-FP-8211** | male | stage iib | Cardia, NOS | nonEBV |
| **TCGA-FP-8631** | male | stage iiia | Cardia, NOS | nonEBV |
| **TCGA-FP-A4BF** | male | stage iiia | Cardia, NOS | nonEBV |
| **TCGA-FP-A8CX** | male | stage iiic | Body of stomach | nonEBV |
| **TCGA-FP-A9TM** | male | not reported | Cardia, NOS | nonEBV |
| **TCGA-HF-7132** | male | stage ii | Gastric antrum | nonEBV |
| **TCGA-HF-7133** | female | stage iv | Stomach, NOS | nonEBV |
| **TCGA-HF-7136** | male | stage iiia | Cardia, NOS | nonEBV |
| **TCGA-HF-A5NB** | female | stage iiic | Gastric antrum | nonEBV |
| **TCGA-HU-8249** | male | stage iiia | Body of stomach | nonEBV |
| **TCGA-HU-8604** | female | stage iia | Gastric antrum | nonEBV |
| **TCGA-HU-8608** | male | stage iiib | Body of stomach | EBV |
| **TCGA-HU-8610** | male | stage ia | Body of stomach | nonEBV |
| **TCGA-HU-A4G2** | male | stage iib | Gastric antrum | EBV |
| **TCGA-HU-A4G3** | male | stage iib | Gastric antrum | nonEBV |
| **TCGA-HU-A4G6** | male | stage ia | Cardia, NOS | EBV |
| **TCGA-HU-A4G8** | female | stage iib | Gastric antrum | nonEBV |
| **TCGA-HU-A4G9** | female | stage ia | Body of stomach | nonEBV |
| **TCGA-HU-A4GC** | male | stage iiib | Gastric antrum | nonEBV |
| **TCGA-HU-A4GD** | male | stage iib | Gastric antrum | nonEBV |
| **TCGA-HU-A4GF** | male | stage iia | Gastric antrum | nonEBV |
| **TCGA-HU-A4GH** | male | stage ia | Body of stomach | nonEBV |
| **TCGA-HU-A4GN** | male | stage iia | Gastric antrum | nonEBV |

| | | | | |
|---|---|---|---|---|
| **TCGA-HU-A4GP** | female | stage iia | Gastric antrum | nonEBV |
| **TCGA-HU-A4GQ** | male | stage iiic | Gastric antrum | nonEBV |
| **TCGA-HU-A4GT** | female | stage iia | Gastric antrum | nonEBV |
| **TCGA-HU-A4GU** | male | stage iib | Gastric antrum | nonEBV |
| **TCGA-HU-A4GX** | female | stage iiic | Gastric antrum | nonEBV |
| **TCGA-HU-A4GY** | female | stage iiia | Gastric antrum | nonEBV |
| **TCGA-HU-A4H0** | male | stage iiic | Body of stomach | EBV |
| **TCGA-HU-A4H5** | male | stage ib | Fundus of stomach | nonEBV |
| **TCGA-HU-A4H6** | female | stage iiia | Gastric antrum | nonEBV |
| **TCGA-HU-A4H8** | male | stage ib | Gastric antrum | nonEBV |
| **TCGA-HU-A4HD** | male | stage iiia | Gastric antrum | nonEBV |
| **TCGA-IN-7806** | male | stage iib | Cardia, NOS | nonEBV |
| **TCGA-IN-7808** | male | not reported | Cardia, NOS | nonEBV |
| **TCGA-IP-7968** | male | stage iiib | Cardia, NOS | nonEBV |
| **TCGA-KB-A6F7** | female | stage ib | Body of stomach | nonEBV |
| **TCGA-KB-A93J** | male | stage ii | Cardia, NOS | nonEBV |
| **TCGA-MX-A5UG** | male | stage iiia | Gastric antrum | nonEBV |
| **TCGA-MX-A5UJ** | female | stage iiia | Gastric antrum | nonEBV |
| **TCGA-MX-A663** | male | stage iia | Stomach, NOS | nonEBV |
| **TCGA-MX-A666** | male | stage iia | Stomach, NOS | nonEBV |
| **TCGA-R5-A7O7** | male | stage iv | Cardia, NOS | nonEBV |
| **TCGA-R5-A7ZE** | female | stage iiia | Cardia, NOS | nonEBV |
| **TCGA-R5-A7ZF** | female | stage iv | Gastric antrum | nonEBV |
| **TCGA-R5-A7ZI** | female | stage iv | Stomach, NOS | nonEBV |
| **TCGA-R5-A7ZR** | female | stage iii | Body of stomach | nonEBV |
| **TCGA-R5-A805** | male | stage iiib | Cardia, NOS | nonEBV |
| **TCGA-RD-A7BS** | male | stage iiia | Cardia, NOS | nonEBV |

| | | | | |
|---|---|---|---|---|
| **TCGA-RD-A7BW** | female | stage ib | Fundus of stomach | nonEBV |
| **TCGA-RD-A7C1** | male | stage ib | Fundus of stomach | nonEBV |
| **TCGA-RD-A8MV** | male | stage iiib | Body of stomach | nonEBV |
| **TCGA-RD-A8MW** | male | stage iiib | Gastric antrum | nonEBV |
| **TCGA-RD-A8N0** | female | stage iiib | Fundus of stomach | nonEBV |
| **TCGA-RD-A8N1** | male | stage iiib | Body of stomach | nonEBV |
| **TCGA-RD-A8N4** | female | stage iiia | Gastric antrum | nonEBV |
| **TCGA-RD-A8N6** | female | stage iiia | Gastric antrum | nonEBV |
| **TCGA-RD-A8N9** | female | stage ii | Cardia, NOS | nonEBV |
| **TCGA-RD-A8NB** | female | stage iiia | Cardia, NOS | nonEBV |
| **TCGA-SW-A7EA** | female | stage ib | Gastric antrum | nonEBV |
| **TCGA-SW-A7EB** | male | stage iiia | Gastric antrum | nonEBV |
| **TCGA-VQ-A8DT** | male | stage iiib | Body of stomach | nonEBV |
| **TCGA-VQ-A8DU** | male | stage iiia | Gastric antrum | nonEBV |
| **TCGA-VQ-A8DV** | male | stage ib | Gastric antrum | nonEBV |
| **TCGA-VQ-A8DZ** | male | stage iv | Gastric antrum | nonEBV |
| **TCGA-VQ-A8E0** | male | stage iiia | Body of stomach | nonEBV |
| **TCGA-VQ-A8E2** | male | stage iiib | Body of stomach | nonEBV |
| **TCGA-VQ-A8E3** | male | stage iia | Gastric antrum | nonEBV |
| **TCGA-VQ-A8E7** | male | stage iv | Cardia, NOS | nonEBV |
| **TCGA-VQ-A8P2** | male | stage iiia | Body of stomach | nonEBV |
| **TCGA-VQ-A8P3** | male | stage iiia | Body of stomach | nonEBV |
| **TCGA-VQ-A8P5** | male | stage iia | Body of stomach | nonEBV |
| **TCGA-VQ-A8P8** | female | stage iib | Body of stomach | nonEBV |
| **TCGA-VQ-A8PB** | female | stage ii | Gastric antrum | nonEBV |
| **TCGA-VQ-A8PC** | male | stage iiia | Gastric antrum | nonEBV |
| **TCGA-VQ-A8PD** | male | stage iiic | Body of stomach | nonEBV |

| | | | | |
|---|---|---|---|---|
| **TCGA-VQ-A8PE** | male | stage iiib | Cardia, NOS | nonEBV |
| **TCGA-VQ-A8PF** | male | stage iiib | Body of stomach | EBV |
| **TCGA-VQ-A8PH** | male | stage iiib | Gastric antrum | nonEBV |
| **TCGA-VQ-A8PJ** | male | stage iv | Body of stomach | nonEBV |
| **TCGA-VQ-A8PK** | male | stage iiib | Cardia, NOS | nonEBV |
| **TCGA-VQ-A8PM** | male | stage iv | Cardia, NOS | nonEBV |
| **TCGA-VQ-A8PO** | male | stage iib | Gastric antrum | nonEBV |
| **TCGA-VQ-A8PP** | male | stage iv | Body of stomach | nonEBV |
| **TCGA-VQ-A8PQ** | female | stage iv | Cardia, NOS | nonEBV |
| **TCGA-VQ-A8PU** | female | stage iiia | Body of stomach | nonEBV |
| **TCGA-VQ-A8PX** | male | stage ia | Stomach, NOS | nonEBV |
| **TCGA-VQ-A91A** | male | stage iiib | Body of stomach | nonEBV |
| **TCGA-VQ-A91D** | male | stage iiic | Body of stomach | nonEBV |
| **TCGA-VQ-A91K** | male | stage iiia | Gastric antrum | nonEBV |
| **TCGA-VQ-A91N** | female | stage iv | Gastric antrum | nonEBV |
| **TCGA-VQ-A91Q** | male | stage iv | Cardia, NOS | nonEBV |
| **TCGA-VQ-A91S** | male | stage iiib | Body of stomach | nonEBV |
| **TCGA-VQ-A91U** | male | stage iiia | Cardia, NOS | nonEBV |
| **TCGA-VQ-A91V** | male | stage iiia | Cardia, NOS | nonEBV |
| **TCGA-VQ-A91W** | male | stage iiia | Body of stomach | EBV |
| **TCGA-VQ-A91X** | male | stage iiib | Gastric antrum | nonEBV |
| **TCGA-VQ-A91Y** | male | stage iiic | Gastric antrum | nonEBV |
| **TCGA-VQ-A91Z** | female | stage iiia | Gastric antrum | nonEBV |
| **TCGA-VQ-A922** | male | stage iv | Gastric antrum | nonEBV |
| **TCGA-VQ-A923** | male | stage iiia | Body of stomach | EBV |
| **TCGA-VQ-A924** | male | stage ii | Gastric antrum | nonEBV |
| **TCGA-VQ-A925** | male | stage iiia | Gastric antrum | nonEBV |

| | | | | |
|---|---|---|---|---|
| **TCGA-VQ-A927** | male | stage iiib | Body of stomach | nonEBV |
| **TCGA-VQ-A928** | male | stage iv | Body of stomach | nonEBV |
| **TCGA-VQ-A92D** | male | stage ib | Cardia, NOS | nonEBV |
| **TCGA-VQ-A94O** | male | stage iiic | Gastric antrum | nonEBV |
| **TCGA-VQ-A94P** | male | not reported | Body of stomach | nonEBV |
| **TCGA-VQ-A94R** | male | not reported | Gastric antrum | nonEBV |
| **TCGA-VQ-A94T** | male | stage iiib | Cardia, NOS | nonEBV |
| **TCGA-VQ-A94U** | male | stage iib | Gastric antrum | nonEBV |
| **TCGA-VQ-AA64** | male | stage iiib | Cardia, NOS | nonEBV |
| **TCGA-VQ-AA68** | female | stage iiic | Cardia, NOS | nonEBV |
| **TCGA-VQ-AA69** | male | stage iiia | Cardia, NOS | EBV |
| **TCGA-VQ-AA6A** | male | stage iiic | Cardia, NOS | nonEBV |
| **TCGA-VQ-AA6B** | male | stage iiib | Cardia, NOS | nonEBV |
| **TCGA-VQ-AA6D** | female | stage iiia | Gastric antrum | nonEBV |
| **TCGA-VQ-AA6F** | male | stage iib | Cardia, NOS | nonEBV |
| **TCGA-VQ-AA6G** | male | stage iia | Cardia, NOS | nonEBV |
| **TCGA-VQ-AA6I** | male | stage iiib | Cardia, NOS | nonEBV |
| **TCGA-VQ-AA6J** | male | stage iiib | Cardia, NOS | nonEBV |
| **TCGA-VQ-AA6K** | male | stage iiic | Cardia, NOS | nonEBV |
| **TCGA-ZA-A8F6** | male | stage ib | Gastric antrum | nonEBV |
| **TCGA-ZQ-A9CR** | female | stage iiic | Gastric antrum | nonEBV |

**Suppl. Table 3: Clinical characterization of TCGA-STAD (N=317 patients).** NOS = not otherwise specified.
